# Position-independent functional refinement within the vagus motor topographic map

**DOI:** 10.1101/2023.09.11.557289

**Authors:** Takuya Kaneko, Jonathan Boulanger-Weill, Adam J Isabella, Cecilia B Moens

**Affiliations:** Division of Basic Sciences, Fred Hutchinson Cancer Center, Seattle, Washington 98109, USA; Department of Molecular and Cellular Biology, Faculty of Arts and Sciences, Harvard University, Cambridge, MA 02138, USA; Sorbonne Université, Institut National de la Santé et de la Recherche Médicale, Centre National de la Recherche Scientifique, Institut de la Vision, Paris, France; Department of Genetics, Cell Biology, and Development, University of Minnesota, Minneapolis, Minnesota 55455, USA

## Abstract

Motor neurons in the central nervous system often lie in a continuous topographic map, where neurons that innervate different body parts are spatially intermingled. This is the case for the efferent neurons of the vagus nerve, which innervate diverse muscle and organ targets in the head and viscera for brain-body communication. It remains elusive how neighboring motor neurons with different fixed peripheral axon targets develop the separate somatodendritic (input) connectivity they need to generate spatially precise body control. Here we show that vagus motor neurons in the zebrafish indeed generate spatially appropriate peripheral responses to focal sensory stimulation even when they are transplanted into ectopic positions within the topographic map, indicating that circuit refinement occurs after the establishment of coarse topography. Refinement depends on motor neuron synaptic transmission, suggesting that an experience-dependent periphery-to-brain feedback mechanism establishes specific input connectivity amongst intermingled motor populations.

## INTRODUCTION

Both sensory and motor neurons form topographic maps in which the positions of neurons in the central nervous system (CNS) reflect the spatial organization of the body parts they innervate^1–4^. Topography in motor neurons is thought to enable appropriate connectivity to upstream neurons^5,6^. In proprioceptive circuits in the spinal cord, for example, loss of motor pool topographic organization results in inappropriate wiring to upstream sensory neurons and consequently, abnormal motor behavior^7^. Unlike in the spinal cord, motor maps in the brain are continuous, with the cell bodies and dendrites of neurons responsible for one motor response loosely distributed and intermingled with neurons that control other responses^4,8–12^. Shared somatodendritic (input) connectivity amongst motor neurons in the same topographic position would thus potentially lead to concurrent activation of neurons that elicit different body functions. It remains poorly understood whether neighboring neurons in a continuous motor map differ in input connectivity and, if so, how neurons develop specific input connectivity that is appropriate for their axonal targeting.

In this study, we address this question by investigating motor efferents of the vagus nerve (the 10^th^ cranial nerve, nX) which mediate brain-body communication by innervating a wide array of body parts including muscles of the pharynx and larynx and post-synaptic parasympathetic neurons in most visceral organs. These diverse outputs control coughing, gagging, swallowing, heart rate, stomach emptying, esophagus and gut peristalsis, and pancreas and gall bladder secretions^13–16^. Despite their functional diversity, all the vagus motor neurons (mXns) lie in two motor nuclei in the brainstem in mammals (the dorsal nucleus, DMV and the nucleus ambiguous, nAmb) or a single vagus motor nucleus in fish^17,18^ where they are organized into a continuous topographic map^8,10,19–21^.

Within the vagus motor topographic map there is significant spatial overlap between neurons that innervate different muscles in the pharynx, which lack any known molecular distinctions^11,19^. In mammals, pharynx-innervating mXns in the nAmb are further intermingled with mXns that innervate the heart and lung^22^. Imprecise wiring of pharynx-innervating motor neurons to upstream circuits can have adverse consequences such as dysphagia (defective swallowing), situational syncope (fainting triggered by swallowing or coughing) and, in more severe cases, sudden cardiac or respiratory arrest during swallowing or gagging as implicated in sudden infant death syndrome^23–27^. The mechanism by which intermingled mXns are wired for precise body control is unknown.

Here we establish the larval zebrafish vagus nerve as a tractable model for studying the fine-scale wiring of intermingled motor neuron groups. First, we demonstrate that pharynx-innervating mXns can deliver spatially precise pharyngeal control through differential activation of neighboring mXns, supporting fine-scale connectivity precision amongst intermingled mXns. We provide evidence that pharyngeal control is refined after the mXn topographic map is formed and axon targeting is fixed, and that this refinement can occur independently of mXn position in the topographic map. We further show that input connectivity of motor neurons depends on the synaptic output from the motor axon. This study suggests a novel experience-dependent periphery-to-brain feedback that modifies motor connectivity of single neurons to provide specific input connectivity within a continuous topographic map.

## RESULTS

### Vagus motor neuron target groups are intermingled in the zebrafish brainstem

After projecting from the brainstem as a single fascicle, the vagus motor nerve in zebrafish splits into five major branches in the periphery (Fig. 1A, B)^8,21^. The first three branches innervate muscles of pharyngeal arches (PA) 4-6 for swallowing and gill opening and closing (branches 4-6) (Fig. 1A). We refer to the mXns that project in these three branches as pharynx-innervating mXns. In some fish, pharynx-innervating mXns control a complex intra-oral food sorting behavior that separates and regurgitates unpalatable materials from edible ones through a 3-neuron reflex circuit (sensory>interneuron>motor)^28,29^. The fourth vagus branch innervates the chewing muscles of the tooth-bearing pharyngeal arch 7, and the fifth branch projects towards the viscera where it branches further to innervate separate organs (branch v in Fig. 1A). An individual mXn only selects a single branch, and once an axon selects a branch it does not change^8,21^. We refer to the group of mXns that contribute to an individual branch of the vagus nerve as a "target group”.

**Figure 1.**
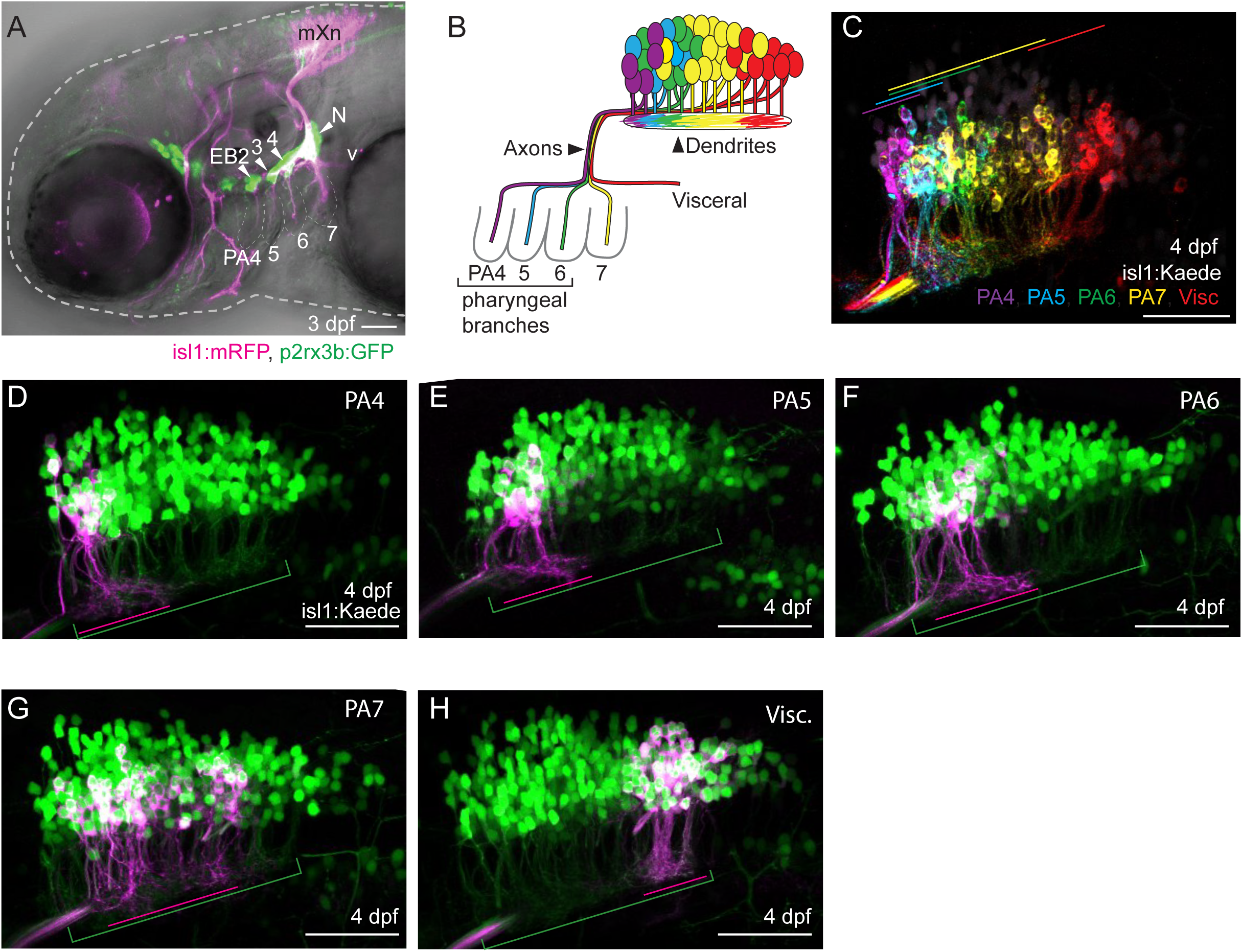
Vagus motor neuron target groups are intermingled in the brain. **(A)** 3 day post-fertilization (dpf) larva with cranial motor neurons in magenta and cranial sensory neurons in green. Vagus motor neurons (mXn) innervate the pharyngeal arches (PA4-7) and the viscera (v). Vagus sensory neurons lie in epibranchial ganglia (EB2-4) associated with PA4-6 respectively, and in the large nodose ganglion (N). **(B)** Schematic of mXn topography. Vagus motor neurons that target different branches (“target groups”) are shown in different colors. Reproduced from Isabella AJ et al. 2020 ^21^. **(C)** mXn target groups in a single 4 dpf larva visualized by sequential retrograde labeling and imaging. Each target group is shown in a different color, with corresponding colored lines indicating the anterior-posterior (A-P) extent of each target group territory. **(D-H)** Retrograde labeling of individual target groups in 4 dpf larvae showing dendritic field overlap. Lines indicate the A-P extent of the individual target group dendrites (magenta) and the entire mXn dendritic field (green). For overlap quantitation see Fig. S1. Scale bars: 50 µm.

The cell bodies of the five target groups are topographically organized along the anterior-posterior (A-P) extent of the nucleus, with progressively more posterior mXns innervating progressively more posterior branches (Fig. 1B)^8,21^. However, within this branch-level topography, mXns are continuously mapped, with no separation between target groups (Fig.1B, C). In particular, the three target groups constituting pharynx-innervating branches PA4-6, visualized by the sequential photoconversion and imaging of individual branches in a single Tg(*isl1:Kaede*)^30^ fish, are highly intermingled within the anterior part of the nucleus, so that mXns from different target groups are frequently positioned at the same A-P level (Fig. 1C-F). Likewise, viscera-targeting mXns occupy the posterior end of the nucleus; but there is no apparent separation into distinct domains for mXns that innervate different organs, analogous to the continuous topography of the mammalian DMV (Fig. 1C, H)^20^. Lastly, mXns that innervate branch 7 are widely scattered along the A-P extent of the nucleus (Fig 1C, G), overlapping with both pharynx-innervating mXns anteriorly and viscera-innervating mXns posteriorly.

Motor dendrites, which develop after motor axons have selected their peripheral branch targets, extend longitudinally in a neuropil ventral to the motor nucleus. Like the cell bodies from which they extend, mXn dendrites of different target groups have an overlapping, continuous topographic organization (Fig.1D-H). The spatial overlap of target groups is particularly evident amongst the dendrites of pharynx-innervating mXns (Fig. 1C-F, S1). These observations are consistent with reports of significant spatial overlap between mammalian mXns that innervate different pharyngeal muscles^11^ and the lack of unique dendritic morphology amongst neighboring mXns^19,31^, and therefore predict a conserved wiring strategy across vertebrates.

### Focal noxious stimulation to the pharynx elicits motor responses that become more specific over time

To determine whether different pharyngeal muscles can be differently controlled despite the spatial overlap between the mXns that innervate them, we sought to establish a method to reproducibly induce pharyngeal muscle contractions. We reasoned that noxious stimulation to the pharynx would induce reflexive pharyngeal muscle contraction in zebrafish, analogous to the protective gag reflex which mammals and some fish use to expel inedible materials from the pharynx^28,29^. We delivered noxious stimulation locally to the pharynx with optovin, a photo-activatable ligand for TrpA1 nociceptive channels^32^ and visualized sensory neuron responses using the calcium indicator GCaMP6s in Trpa1b-expressing sensory neurons (Tg(*TrpA1b^Gal^*^4^; *UAS:GCaMP6s*) in 7 days post-fertilization (dpf) larvae. In the presence of optovin, focal UV illumination of the anterior pharynx (between PA3 and PA4) and posterior pharynx (between PA6 and PA7) activates a largely non-overlapping set of Trpa1b-expressing vagus sensory neurons in the nodose and epibranchial ganglia (Fig. S2A-D). The central axon terminals of Trpa1b vagus sensory neurons are organized topographically dorsal to the vagus motor dendritic neuropil (Fig. S2A, E): the central processes of anterior PA-innervating sensory neurons lie anterior to those of posterior PA-innervating sensory neurons, though with significant spatial overlap (Fig. S2E, F). Consistent with this organization, the activity patterns in the central axons based on calcium imaging are topographically organized with anterior stimulation exciting anterior processes and posterior stimulation exciting posterior processes (Fig. S2G-I).

We found that focal UV illumination to the pharynx in the presence of optovin reproducibly induces immediate muscle contraction as we expected (Movie 1, 2). To compare the level of muscle activity between neighboring pharyngeal muscles, we expressed GCaMP6s in the muscles with *tcf21^Gal4^;* Tg(*UAS:GCaMP6s*) (Fig. 2A)^33^. We observed that although focal noxious stimulation often induces GCaMP responses from all of the PA4, PA5, and PA6 muscles, the relative levels are different. Focal noxious stimulation of the anterior pharynx induces a stronger response in the PA4 muscle, near the stimulation spot, than in the PA5 and PA6 muscles (Fig. 2B, C). Conversely, focal stimulation of the posterior pharynx induces a stronger response in the nearby PA5 and PA6 muscles than the more distant PA4 muscle. The muscle response is dependent on the presence of optovin (Fig. 2D). Strong responses by muscles near the stimulation spots together with weaker responses by more distant muscles may generate the coordinated pharyngeal constriction that effectively expel unpalatable objects from the pharynx.

**Figure 2.**
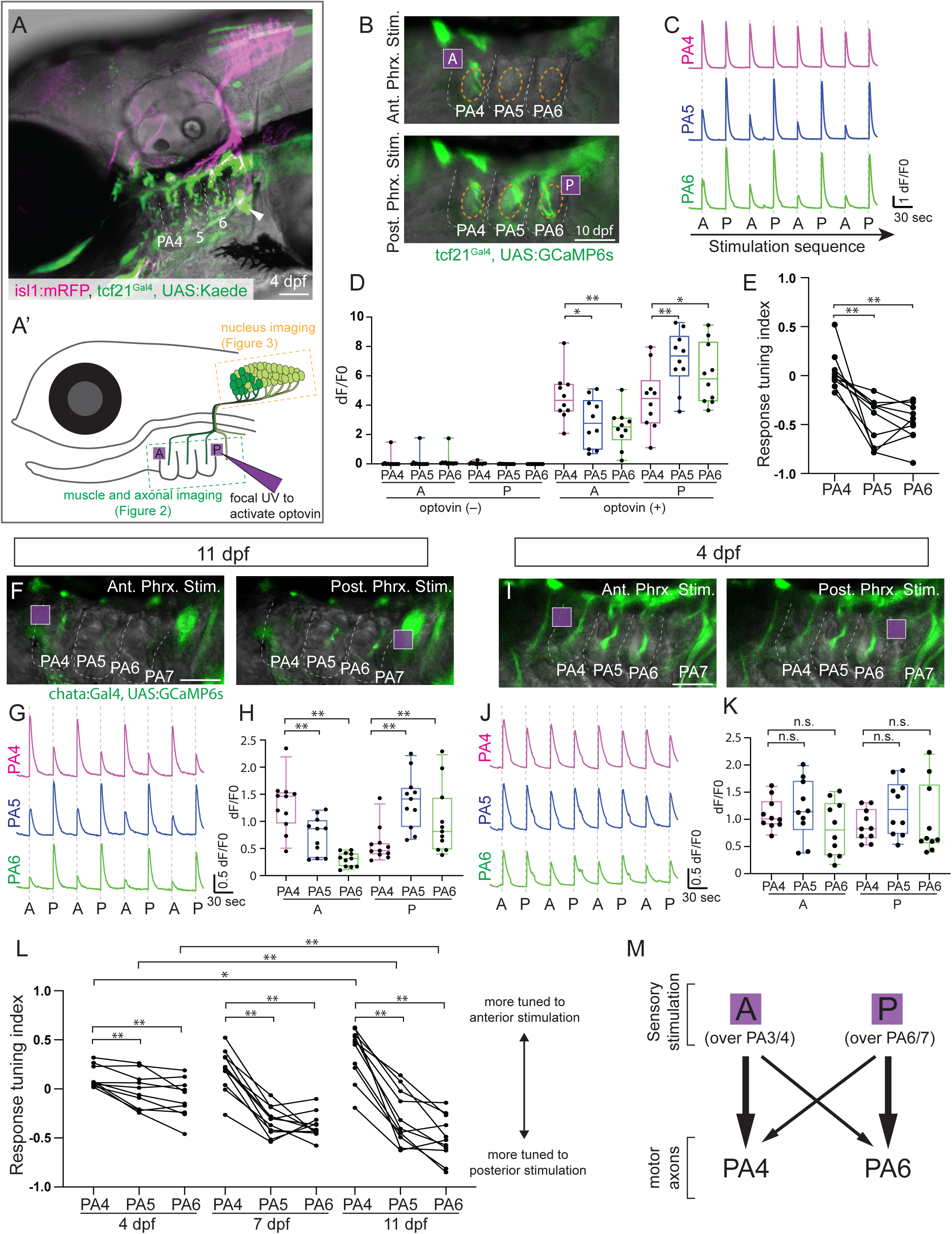
Focal noxious stimulation to the pharynx elicits motor responses that become more specific over time. **(A)** 4 dpf fish showing motor neurons (magenta) and cranial muscles (green). The PA4, PA5, and PA6 branches innervate structurally equivalent muscles while the PA7 branch innervates a different muscle (arrowhead). **(A’)** Schematic showing the anterior (A) and posterior (P) pharyngeal spots illuminated by UV laser for optovin activation. The green and orange rectangles show the calcium imaging area shown in Fig. 2 and Fig. 3, respectively. **(B)** GCaMP6s activity in pharyngeal muscles in response to A (top) and P (bottom) stimulation at 10 dpf. The regions of interest (ROIs) used for quantification are indicated by orange circles. **(C)** mean calcium traces for pharyngeal muscle responses from 10 animals over 4 iterations of alternating A and P stimulation. **(D)** Comparison of mean calcium responses between PA4, PA5 and PA6 muscles in the absence and presence of optovin at 10 dpf. n=10 larvae for each condition. **(E)** Tuning of the calcium response of PA4, PA5 and PA6 muscles to focal stimulation at 10 dpf (see text for explanation of the “response tuning index”). **(F, I)** GCaMP6s activity in vagus motor branches innervating PA4-6 at 11 dpf (**F**) and 4 dpf (**I**) in response to A (left) and P (right) stimulation. **(G, J)** Mean calcium traces of motor branch responses over 4 iterations of alternating A and P stimulation at 11 dpf (**G**, n=11) and 4 dpf (**J**, n=10). **(H, K)** Comparison of calcium responses between PA4, PA5 and PA6 branches at 11 dpf (**H,** n=11) and 4 dpf (**K**, n=10). **(L)** Response tuning index of PA4, PA5 and PA6 mX branches at 4 dpf (n=10), 7 dpf (n=11), and 11 dpf (n=11). **(M)** Diagram showing the relationship between stimulation locations and responding motor branches. Scale bar 50 µm. Statistics in panels **D**, **H** and **K**: paired t-test. Statistics in panels **E** and **L**: Wilcoxon matched-pairs singled rank test. ** indicates p<0.01, * indicates 0.01<p<0.05, n.s. indicates 0.05<p throughout the paper.

We quantified the difference in calcium responses to the two pharyngeal stimulations as “tuning” in which an exclusive response to anterior pharynx stimulus would give a "tuning index” score of 1 and exclusive response to a posterior stimulus a score of −1 (see methods). The anterior PA4 muscle and the posterior PA5, PA6 muscles exhibit tuning differences which are stereotypic across animals (Fig. 2E). Taken together, in spite of extensive topographic overlap, pharynx-innervating mXns are able to control individual pharyngeal muscles differently, generating stereotypic muscle contraction patterns along the A-P axis, corresponding to the position of the noxious stimulus.

Next we directly examined whether pharynx-innervating mXns, upstream of pharyngeal muscles, show spatially regulated activity patterns. We imaged GCaMP6s signals in the motor axon branches innervating PA4, 5, and 6 in 10-11 dpf Tg(*chata:Gal4*; *UAS:GCaMP6s*) larvae in response to focal noxious stimulation mediated by optovin, as above (Fig. 2F). Chata is expressed in motor neurons and not sensory neurons. As expected from the activity patterns in pharyngeal muscles, we find that anterior pharynx stimulation robustly activates the PA4 branch with weaker activity from PA5 and PA6 branches, while posterior pharynx stimulation strongly activates PA5 and PA6 branches with weaker activity from the PA4 branch (Fig. 2F-H). Each animal shows the same pattern of axonal activities following repeated delivery of sensory cues, and the responses are stereotyped with little variation across different larvae (Fig. 2G, H). This results in a tuning difference among the three axonal branches, which represents functional separation amongst the mXns innervating them (Fig. 2L, right panel). This observation supports the idea that mXn target groups, though intermingled, are preferentially connected with upstream sensory circuits that detect stimuli in the same anatomical region that the mXn target group innervates.

We examined when the functional separation among pharynx-innervating mXns develops. At 4 dpf, shortly before larval zebrafish begin using mXns for eating, focal noxious stimulation delivered as above induces similar GCaMP activity levels in all three branches (PA4-PA6; Fig. 2I-K). A difference in tuning among pharynx-innervating mXns is still detectable, but it is immature compared to that in older larvae (Fig. 2L, left vs right panel). At 7dpf, tuning differences are more evident but still less mature than that at 10-11 dpf (Fig. 2L, middle panel). Given that the topographic organization of motor neurons that we have described is already established at 3 dpf (Fig 1A, B)^8,21^, and mXn axon targeting does not change after that stage, our data support a model where mXns progressively refine input connectivity for more sophisticated peripheral control after initial motor circuits are formed.

### Focal noxious stimulation elicits responses in spatially overlapping motor neurons

To map the topographic distribution of mXn cell bodies that respond to focal noxious stimulation, we expressed nuclear GCaMP6s in motor neurons (Tg(*isl1:H2B-GCaMP6s*)) and imaged a single focal plane of the vagus motor nucleus during noxious stimulation (Fig. 3A). In addition to anterior and posterior pharynx stimulation, we provided a third more posterior noxious stimulation to the esophagus. In 10-11 dpf larvae, both anterior and posterior pharynx stimulation activates a subset of mXns in the anterior 45 µm of the ∼150 µm-long motor nucleus where pharynx-innervating mXns reside, while esophageal stimulation strongly activates mXns in the posterior-most part of the motor nucleus where viscera-innervating mXns reside (>90 µm)(Fig. 3A-D). Esophageal stimulation occasionally weakly activates some anterior mXns and pharyngeal stimulation weakly activates posterior mXns (Fig. 3B, D). However, pharyngeal stimulation rarely gives high activities from the middle of the motor nucleus (45-90 µm), which are largely occupied by PA7-targeting mXns, suggesting a specific recruitment of mXns responsible for muscle contraction around the location of noxious stimulation.

**Figure 3.**
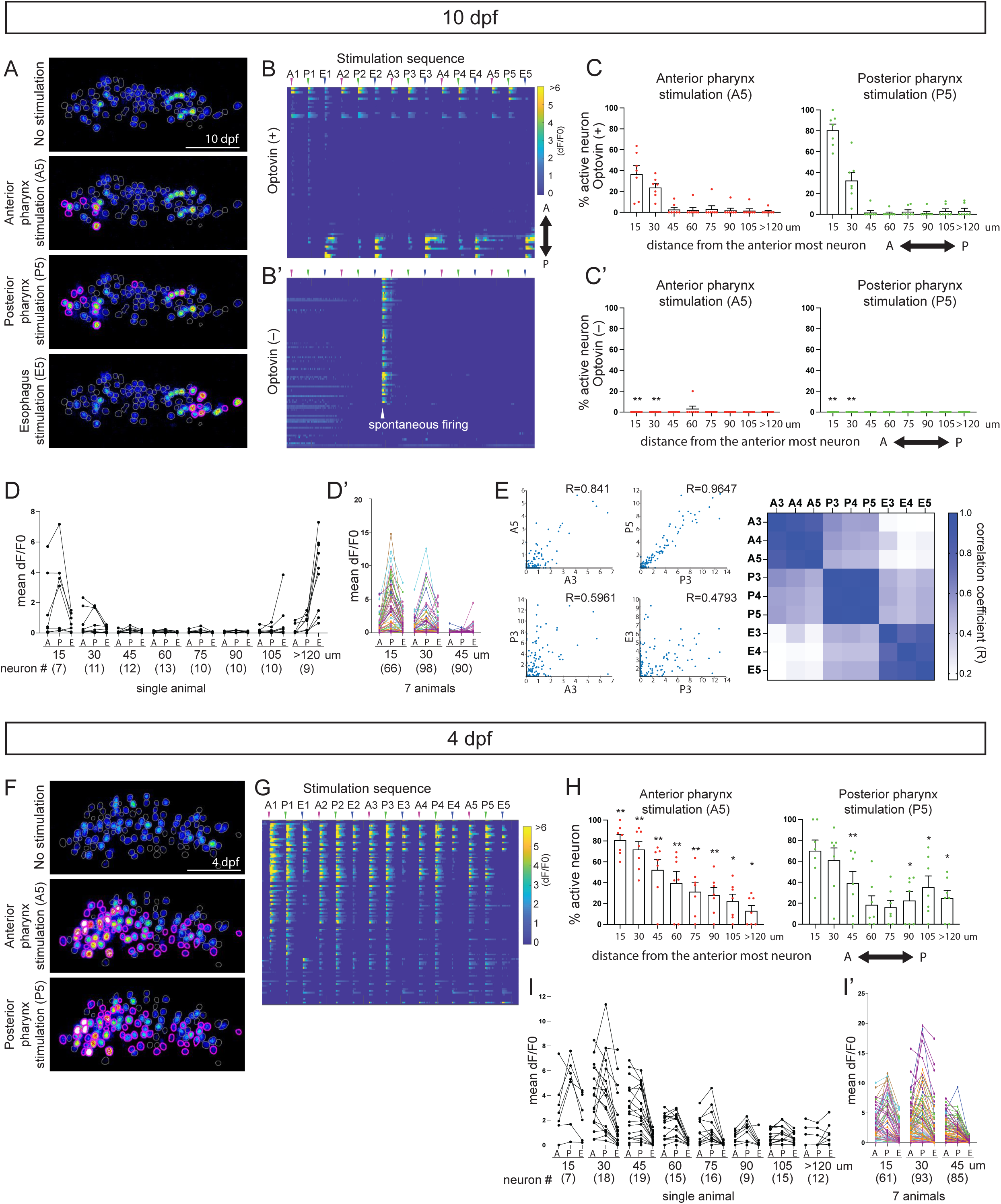
Focal noxious stimulation elicits responses in spatially overlapping motor neurons. **(A, F)** Calcium responses of mXns to anterior pharynx (top), posterior pharynx (middle) and esophagus (bottom) in representative 10 dpf (**A**) and 4 dpf (**F**) larvae. GCaMP6s signals are shown in the heat-color. ROIs assigned by automated segmentation are shown in gray. Magenta ROIs indicate active neurons determined by automated analysis. **(B, G)** Calcium traces for the neurons in the larvae in **A**, **F**, respectively, in response to five cycles of anterior pharynx (A), posterior pharynx (P) and esophagus (E) noxious stimulation. Neurons are aligned from top to bottom corresponding to their anterior-to-posterior position in the nucleus. The magnitude of GCaMP signals is coded in the heat-color. **(B’)**: responses to focal UV illumination in the absence of optovin in 10 dpf larvae. Spontaneous firing (arrowhead) was observed independent of illumination timing. **(C, H)** The percentage of GCaMP6s-expressing neurons in each of 15-µm bins along the A-P axis of the vagus motor nucleus that respond to A5 (left) and P5 (right) pharyngeal stimulation at 10 dpf (**C**, n=7 larvae) and 4 dpf (**H**, n=7 larvae). **(C’)** responses in the absence of optovin (n=7 larvae). Statistics in **C’** and **H** shows t-test comparison with **C**. **(D, I)** Mean dF/F0 in neurons in each of 15-µm bins from the single larva in **A** (for **D**) and **F** (for **I**). Results from 7 animals (in different colors) are shown in **D’**, **I’** for the first three bins. **(E)** Correlation analysis of GCaMP6s responses of mXns to stimuli in the same and different locations^64^. **Left:** Dots represent all analyzed neurons collected from 7 animals, which are plotted based on the GCaMP6s levels (dF/F0) following two noxious stimulations at the same (top graphs) or different (bottom graphs) locations. Correlation coefficient (R) is measured as the indication of correlation levels. **Right:** Correlation matrix for 3 iterations, with the level of correlation (R) color coded. Scale bar in **A, F**: 50 µm.

In spite of activating a largely non-overlapping set of sensory neurons (Fig. S2F), anterior and posterior pharynx stimulation generally recruits a surprisingly common set of mXns in the anterior portion of the nucleus (Fig. 3D). However, individual mXns respond to the two stimuli with different activity levels, suggesting a different strength in their upstream connections with the two sensory pathways. This difference presumably contributes to the stereotypic tuning difference we observed between the axonal branches and between pharyngeal muscles (Fig. 2E, L). Meanwhile, other interspersed mXns are unresponsive to either stimulus (Fig. 3A, B, D). The difference in response pattern among neighboring mXns is not due to stochasticity in each response, because repeated stimulation to the same spot in a given fish induces more correlated activity patterns than stimulation to different spots does (Fig.3E). Taken together, our results suggest that precise body control by the vagus is established by fine-scale connection specificity that differently wires neighboring mXns.

As we expected from the immature axonal activity patterns we observed in 4 dpf larvae (Fig. 2L), the motor nucleus displays immature response patterns at this early stage. Pharyngeal stimulation activates a much higher proportion of mXns in the anterior region of the nucleus at 4 dpf compared to 10-11 dpf, the response is much broader, and differences in the magnitude of GCaMP responses to anterior vs posterior pharynx stimulation are smaller (Fig. 3H-I). Thus, between 4 and 10 dpf, the sensory-evoked response in many mXns becomes suppressed while being selectively maintained in a few mXns. Furthermore, differences in the magnitude of the response to spatially distinct stimuli emerge. The response suppression may be mediated by a refinement of excitatory input and/or inhibitory input.

This progressive refinement of motor responses is not due to a corresponding refinement of sensory neuron responses, as we found that activity patterns in Trpa1b sensory axons are already organized topographically in 4 dpf larvae and remain unchanged in 10 dpf larvae (Fig. S2I). Although these experiments do not rule out fine-scale changes in sensory neuron synapse organization over time or changes in the central circuit organization, they are consistent with a model in which the maturation of the motor response between 4 dpf and 10 dpf is driven by progressive refinement in the input connectivity of mXns.

### Spatially appropriate responses of vagus motor neurons can be established independent of their position in the topographic map

Our data so far have demonstrated that spatially precise pharyngeal control is delivered by differential activation of intermingled mXns in 10-11 dpf larvae. These findings support the idea that individual mXns develop specific input connectivity that is appropriate for their innervation targets, independent of their positioning. This is different from the positional strategy proposed for mammalian spinal motor neurons, in which motor neurons with common targets receive sensory input based on their distinct clustered positions in the spinal cord^5,34^. To directly test whether mXns can develop specific input connectivity irrespective of their position in the topographic map, we manipulated mXn positions through transplantation. First, we collected mXns from various positions in the motor nucleus of a Tg(*chata:Gal4*; *UAS:GCaMP6s, isl1:membRFP*) donor embryo and placed them into the anterior region of the vagus motor nucleus of a host embryo (Fig. 4A, B). We performed these transplants at 40 hpf, a stage at which transplanted mXns regenerate axons to, or near, their original targets^35^, so the topography of regenerated neurons was disrupted. At 10 dpf, after axon regeneration, we imaged GCaMP activity in the regenerated axons in response to anterior and posterior pharynx stimulation. Since typically only a fraction of mXns normally respond to optovin stimulation at this stage (Fig. 3C) we increased the likelihood that donor-derived mXns are incorporated into sensory-motor circuits by silencing neurotransmission by host mXns with expression of the botulinum toxin light chain (*isl1:Gal4; UAS:BoTxBLC-GFP*), a specific inhibitor of neurotransmission release (see below for further explanation)^36^.

**Figure 4.**
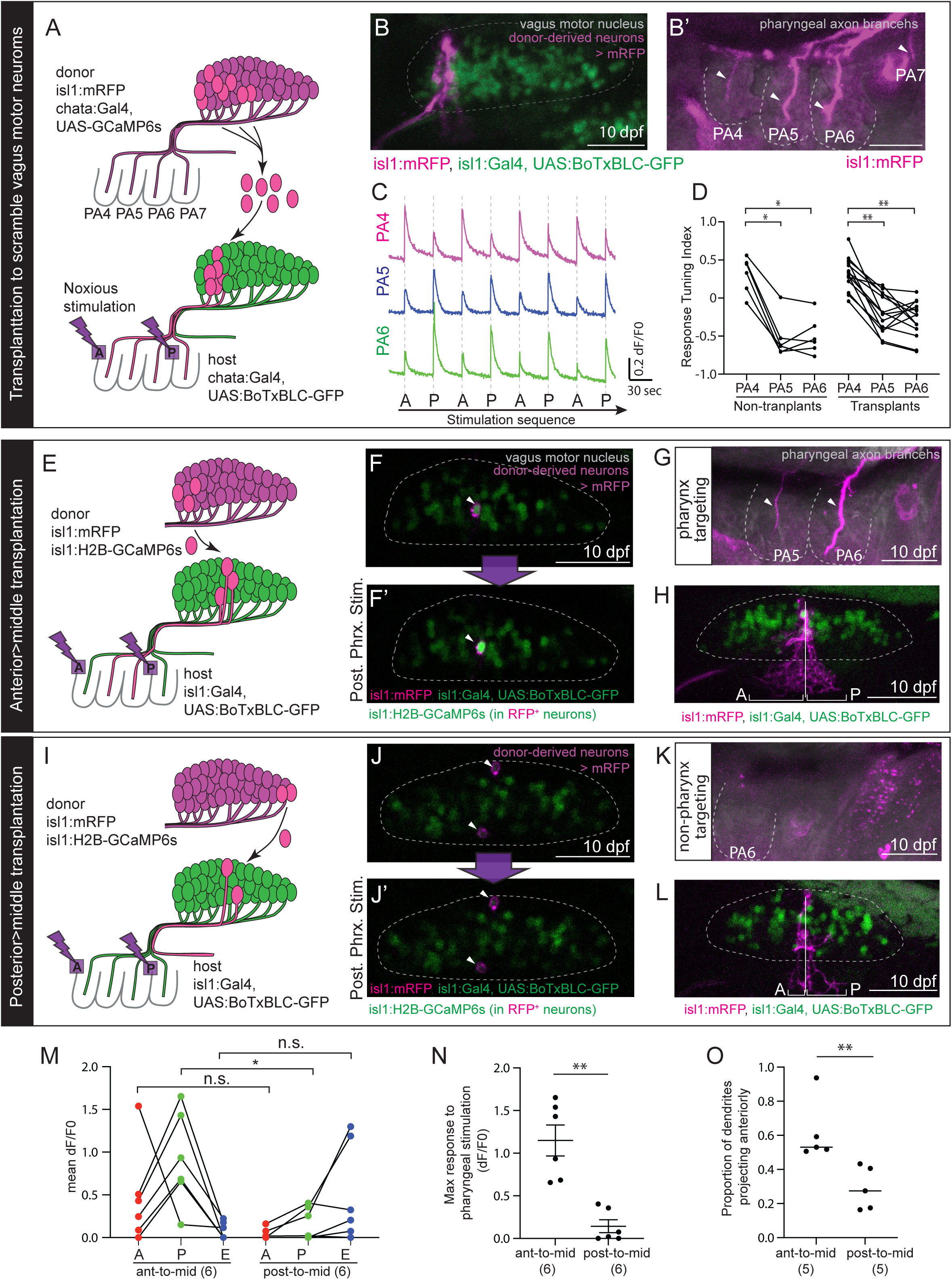
Spatially appropriate responses of vagus motor neurons can be established independent of their position in the topographic map. **(A)** Top: transplantation approach for manipulating the topographic organization of pharynx-targeting mXns. Bottom: at 10 dpf, after axons of transplanted mXns have reinnervated peripheral targets, focal noxious stimulation is delivered as described above to the anterior (A) or posterior (P) pharynx, and GCaMP6s responses in the axons of donor-derived mXns are recorded. GCaMP6s is expressed only in the donor-derived mXns. **(B)** Donor-derived neurons that locate at the same A-P position (magenta) regenerate axons that re-innervate the pharyngeal arches (arrowheads in **B’**). **(C)** Mean GCaMP6s traces of donor-derived axons in response to alternating A and P stimuli from 14 larvae. Signals are weaker than in intact animals (Fig. 2G) because of the smaller number of GCaMP6s-expressing axons and background BoTxBLC-GFP signals. **(D)** Response Tuning Index for n=14 transplanted animals (right) and n=6 untransplanted control animals (left) in which all mXns express GCaMP6s as in Fig. 2G. **(E, I)** Transplantation approaches for generating mispositioned (ectopic) mXns. Anterior (**E**) or posterior (**I**) mXns are transplanted to the middle region of the nucleus (40-60% of its length, a position normally occupied by PA7-innervating mXns that do not respond to either pharynx or esophagus stimulation; see Fig. 3). GCaMP6s responses of transplanted neurons to focal noxious stimulation are measured at 10 dpf. **(F)** GCaMP6s response of mispositioned anterior neuron to posterior pharynx stimulation. **(G, H)** The axons (**G**) and dendrites (**H**) of transplanted anterior-derived mXns in the same larval fish. **(J)** GCaMP6s response of mispositioned posterior neurons to pharynx stimulation. **(K, L)** Absence of axons in the pharyngeal arches (**K**) and posterior polarization of dendrites (**L**) of mispositioned posterior neurons in the same larval fish. **(M)** Mean dF/F0 following noxious stimulation to A, P or esophagus (E) is shown for 6 ectopic anterior-derived (left) and 6 ectopic posterior-derived (right) neurons positioned in the middle of the motor nucleus. **(N)** The higher dF/F0 value from A or P stimulation for each neuron in **M**. **(O)** The proportion of the area of dendrites anterior to the cell body as a fraction of the total dendritic area. 5 animals were used for both conditions. Statistics in **D**: Wilcoxon matched-pairs singled rank test. Statistics in **M, N, O**: t-test. Scale bars 50 µm.

Similar to what we saw in intact fish (Fig. 2G), anterior pharyngeal stimulation preferentially activates donor-derived axons in the PA4 branch, and posterior stimulation robustly activates those in the PA5 and PA6 branches (Fig. 4C). Consequently, the tuning index of pharynx-innervating mXns is similar to that of non-transplanted controls (Fig. 4D). Thus, even when pharynx-innervating mXns are artificially mixed, they can develop specific input connectivity for stereotypical functional outputs, supporting position-independent connection specification.

To further investigate the position-independent strategy, we used heterotopic transplantation to place single GCaMP-expressing mXns away from the other neurons in their target group (Fig. 4E, I). As above, we inhibited neurotransmission from host mXns with BoTxBLC expression. Normally, pharynx optovin stimulation rarely activates endogenous PA7-innervating mXns in the middle of the motor nucleus (Fig. 3C). However ectopic PA4-6-innervating mXns placed in the middle of the motor nucleus frequently respond to pharynx stimulation but not to esophagus stimulation (Fig. 4F, G, M, N). Conversely, ectopic viscera-innervating mXns placed in the middle of the motor nucleus occasionally respond to esophagus stimulation but not to pharynx stimulation (Fig. 4J, K, M, N). Thus, mXns, even if displaced from their normal A-P position and surrounded by mXns of a different target group, can develop the specific input connectivity necessary to generate appropriate motor responses. Our data suggest that mXns can be integrated into correct circuits at the single cell level, independent of neural positioning.

### Ectopic vagus motor neurons form adaptive dendritic projections

What is the mechanism that enables ectopic neurons to integrate into functionally appropriate sensory-motor circuits? By studying the morphology of transplanted mXns after calcium imaging, we noted that transplanted mXns extend dendrites differently depending on where their axons innervate. Whereas the dendritic arbors of posterior-derived mXns preferentially extend posteriorly (Fig. 4L, O), those of anterior-derived pharynx-innervating mXns show relatively more anterior extension (Fig. 4H, O). By extending dendrites towards the neuropil occupied by the dendrites of their target group, transplanted neurons may achieve the correct input connectivity that allows them to respond appropriately to distinct spatial cues.

Supporting this possibility, similar dendritic adjustment was observed when topographically incorrect mXns were generated by an independent method. mXns that express a dominant-negative retinoic acid receptor (DN-RAR) innervate more anterior branches than non-expressing mXns at the same A-P position, due to premature initiation of an axon guidance program^21^. We generated larvae with sparse DN-RAR-expressing neurons (Tg(*isl1:Gal4*; *UAS:DN-RAR-p2A-GFP-CAAX*)). We observed that the dendrites of posteriorly located DN-RAR-expressing mXns, whose axons often innervate the pharynx instead of the viscera, often extend far anteriorly toward the dendritic neuropil of endogenous pharynx-innervating mXns in 7-8 dpf larvae (Fig. 5A-C, E, F). While these adaptive dendritic extensions are more dramatic than would normally be needed for adjacent mXns of different target groups to find the correct input connectivity, it unveils a remarkable ability of mXns to adjust input connectivity so that their axons can generate appropriate motor responses to localized sensory input.

**Figure 5.**
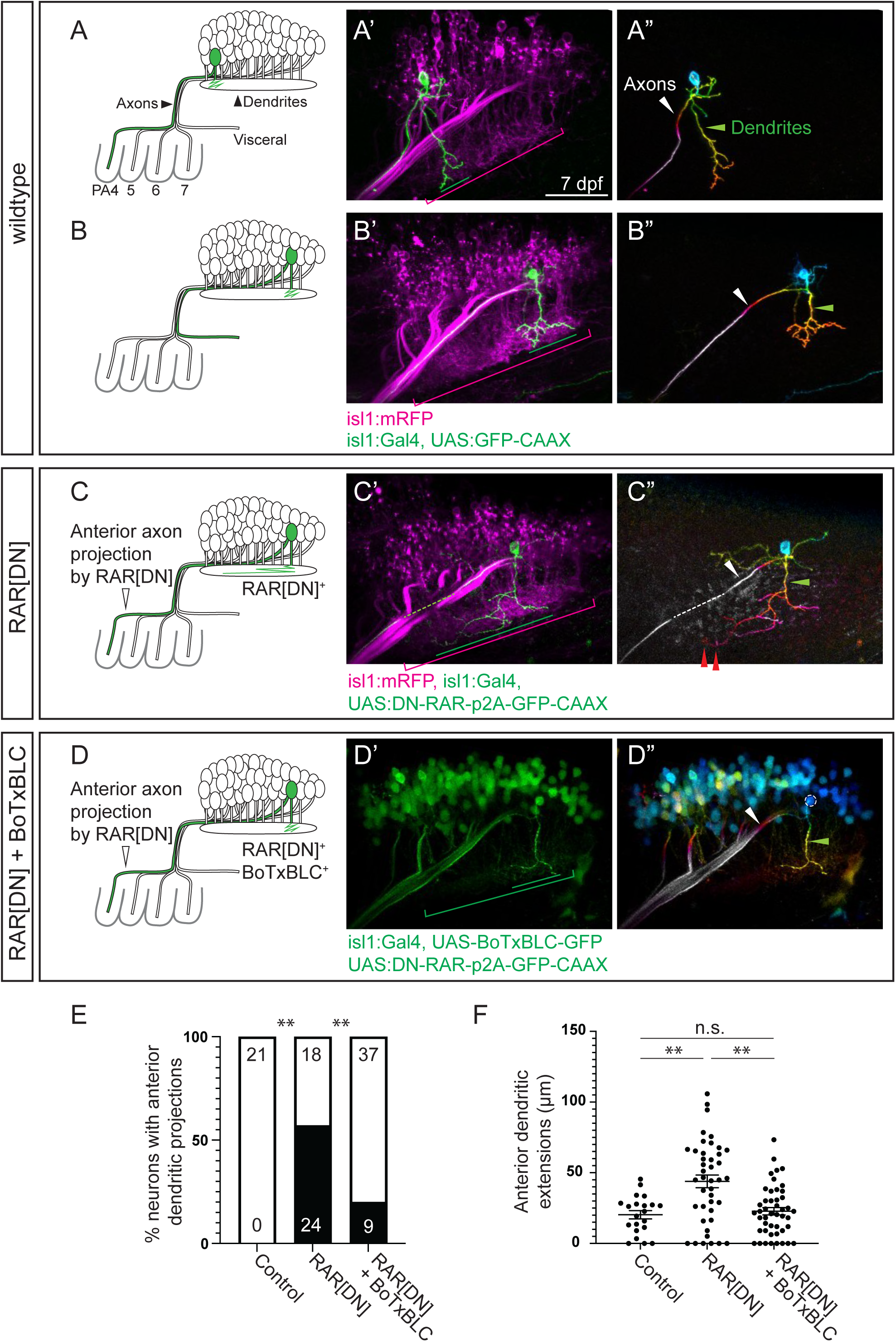
Ectopic vagus motor neurons form adaptive dendritic projections dependent on motor neurotransmission. **(A-C)** Expression of GFP-CAAX (**A, B**; green) or DN-RAR-p2A-GFP-CAAX (**C**; green) in single mXns in the anterior (**A**) or posterior (**B**,**C**) part of the vagus motor nucleus at 7-8 dpf. Green lines in the middle panel indicate the A-P extent of the labeled neuron’s dendrites, and magenta brackets indicate the length of the entire mXn dendritic territory. Red arrowheads in C” indicate the anterior targeted dendrites of a posterior mXn. **(D)** Expression of DN-RAR-p2A-GFP-CAAX (green membrane) in a posterior vagus motor neuron in larva expressing BoTxBLC-GFP (cytoplasmic green) throughout the vagus motor nucleus. **(E)** Comparison of dendritic architecture under the conditions indicated in 7-8 dpf larvae. Posterior mXns whose dendrites extend beyond the posterior 40% of the dendritic neuropil were defined as “neurons with anterior dendritic projections”. The number in the bar graph indicates the sample number used in the analysis. **(F)** Comparison of anterior dendritic extensions (in µm) from the position of the cell body. The same larvae from **E** were analyzed. GFP signals in A’-D’ are depth coded in the heat-color in A”-D”. White arrowheads in A’’-D’’ point to axons; green arrowheads to dendrites. Statistics in **E**: Chi-Square test; in **F**: Mann-Whitney test; scale bar: 50 µm.

### Both neural activity and dendritic adjustment depend on motor neuron output

In many sensory circuits, point-to-point connection specificity is not entirely instructed by genetic programs but established by the experience of sensory inputs^37–39^. Motor circuits can also exhibit experience-dependent synaptic stabilization^40^. In our system, refinement of mXn responses occurs between 4 and 10 dpf, which is after the onset of sensory-evoked activity (Fig. 2L, 3C, H). Thus, we sought to investigate whether refinement of mXn responses depends on mXn activity.

We suppressed the ability of mXns to generate motor outputs by expressing BoTxBLC, which blocks acetylcholine release^36^, using the Gal4/UAS system (Tg(*isl1:Gal4*; *UAS:BoTxBLC-GFP*)). BoTxBLC expression significantly reduces the percentage of anterior mXns that respond to focal pharyngeal stimulation at 10-11 dpf (Fig.6A-D). This effect is not due to defects in motor neuron physiology, because most BoTx-expressing mXns respond strongly to bath application of allyl-isothiocyanate (AITC), a natural agonist of TrpA1 (Fig. 6B-D). Furthermore, we find no change in the topographic organization of cell bodies and dendrites of BoTx-expressing mXns (Fig. 6E, F). Thus, when pharynx-innervating mXns are ineffective in contracting pharyngeal muscles, they become less responsive to pharyngeal noxious cues.

**Figure 6.**
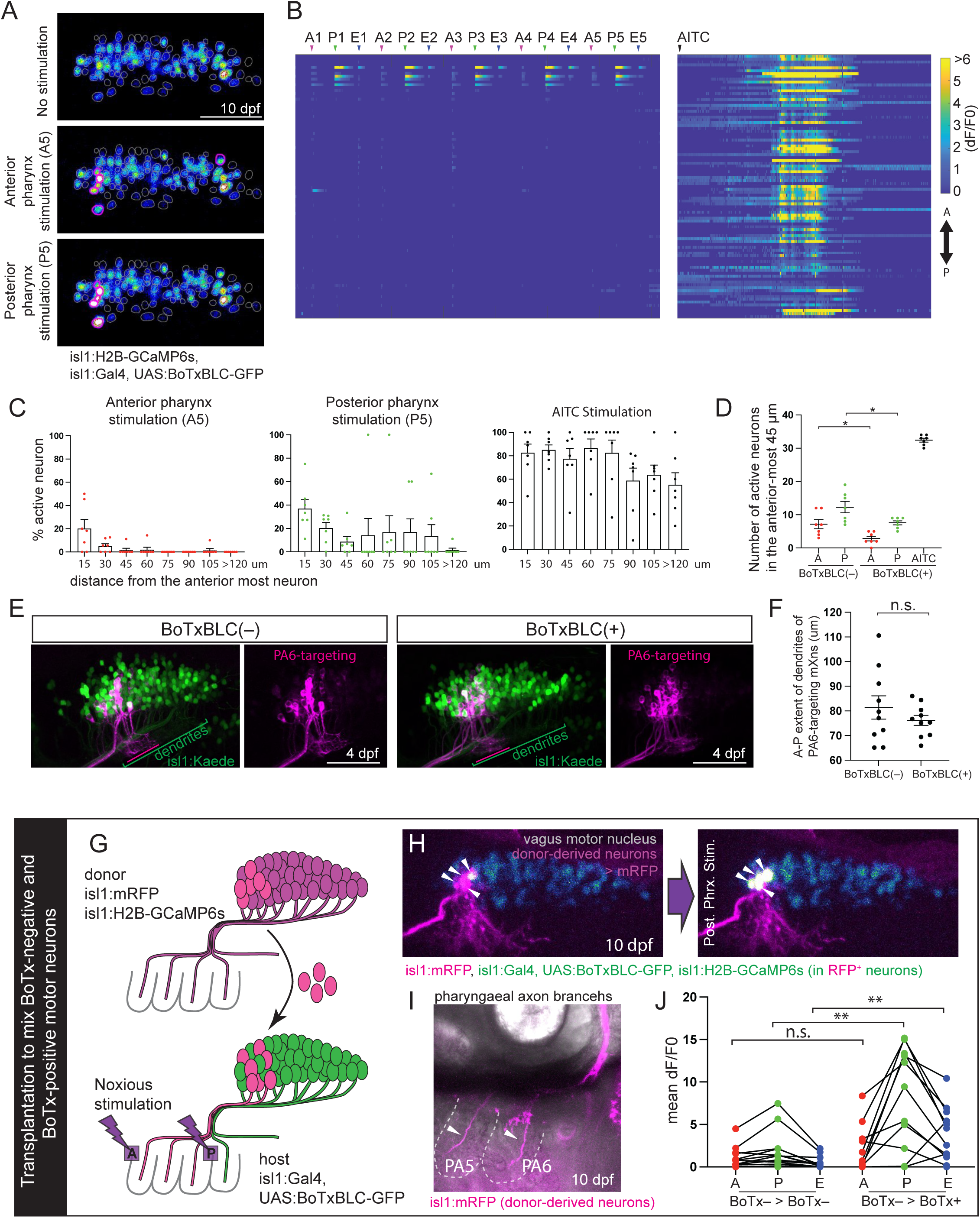
Activity of pharynx-targeting motor neurons depends on their effective neurotransmission. **(A)** Reduced calcium responses in BoTxBLC-expressing mXns to anterior pharynx (A; middle) and posterior pharynx (P; bottom) stimulation in a representative 10 dpf larva (Compare to Fig. 3A) **(B)** Calcium traces for all imaged neurons from the larva in **A** over five cycles of A, P, and E (esophagus) stimulation. Neurons are aligned from top to bottom corresponding to their anterior-to-posterior position in the nucleus. The level of GCaMP signals is coded in the heat map. AITC was applied in the same animal following focal stimulations. **(C)** The percentage of BoTxBLC-expressing mXns in each of 15 µm bins along the A-P length of the motor nucleus that respond to A5 (left), P5 (middle) and AITC stimulation (right) at 10 dpf (n=7 larvae). **(D)** Comparison of the number of mXns responding to the indicated stimulation in the anterior-most 45 µm of the motor nucleus between control (BoTxBLC(-), from Fig. 3C) and BoTxBLC-expressing (BoTxBLC(+)) animals (n=7 larvae per condition). **(E, E’)** Photoconversion of PA6-targeting mXns in control (**E**) and BoTxBLC-expressing (**E’**) larvae at 4dpf. Left panels show unconverted (green) and photoconverted (magenta) neurons while the right panels show only photoconverted neurons. Lines indicate the A-P extent of the photoconverted dendrites (magenta) and the entire mXn dendritic field (green). **(F)** The A-P extent (in µm) of the dendrites of Kaede^Red^+ mXns in **E**. **(G)** Schematic of the transplantation approach for generating BoTx-negative mXns in a BoTx-expressing host vagus motor nucleus. **(H)** Donor-derived mXns (magenta, arrowheads) in a BoTx-expressing host motor nucleus (blue) before (left) and after (right) posterior pharynx stimulation. GCaMP signal (heat-color) in donor-derived mXns (magenta) creates a white pseudocolor. **(I)** Pharyngeal arches of the host larva in **H** innervated by donor-derived axons (magenta). **(J)** Calcium response to A, P and E focal noxious stimulation at 10 dpf of non-BoTx-expressing neurons transplanted into a non-BoTx-expressing (left, n=11) or into a BoTx-expressing (right, n=11) host motor nucleus. Statistics in **D** and **J**: t-test. scale bar: 50 µm.

This effect of BoTxBLC expression is cell-autonomous. When we transplant wildtype (BoTx-negative) mXns into a BoTx-expressing host (Fig. 6G-I)^8,35^, these wildtype neurons are successfully activated by pharyngeal noxious stimulation (Fig. 6H, J). Indeed, they show a significantly higher calcium response than after transplantation into a wildtype host (Fig. 6J). This elevated response in transmission-proficient mXns may reflect a compensatory mechanism that improves peripheral outputs that are otherwise ineffective due to the BoTx-positive background.

Our results together demonstrate experience-dependent refinement, in which inappropriate mXn responses that do not contribute to intended motor outputs are suppressed while appropriate mXn responses are maintained and enhanced as needed. This finding suggests the presence of feedback signaling that reports peripheral motor performance to the brain and accordingly fine-tunes input connectivity. We hypothesize that that this experience-dependent feedback regulation underlies the ability of mXns to adjust their input connectivity to generate accurate motor responses to localized stimuli.

To test this hypothesis, we returned to the sparse DN-RAR expression method to generate posterior mXns that ectopically innervate the pharynx as described above (Fig. 5C), but now blocked neurotransmission by co-expressing BoTxBLC in the entire motor nucleus (Tg(*isl1:Gal4*; *UAS:BoTxBLC-GFP*)). We found that mXns expressing both DN-RAR and BoTxBLC fail to form adaptive dendritic extensions. Rather, they arborize locally at the level of the cell body (Fig. 5D-F) similar to surrounding posterior mXns that project appropriately to the viscera (Fig. 5B, F). Thus, neurotransmission is required for mXns to adapt to position defects by correcting their input connectivity. Taken together, our data support a model in which experience-dependent periphery-to-brain feedback enables individual mXns to detect and fine-tune circuit function.

### Viscera-innervating neurons can integrate into a visceral circuit independent of their position in the motor nucleus

Our study so far has focused on pharynx-innervating mXns due to the substantial spatial overlap amongst target groups in the anterior part of the motor nucleus. We sought to examine whether the position-independent strategy we identified above is a general principle across diverse vagus motor populations. We analyzed viscera-targeting mXns which lie in the posterior part of the motor nucleus.

To specifically induce motor outputs in viscera-innervating mXns, we delivered a mechanosensory stimulus to the esophagus by injecting a droplet of oil into the larval esophagus to mimic a bolus of food (Fig. 7A). Vagus sensory neurons detect esophageal stretch and induce a peristaltic reflex driven by esophagus-innervating mXns to move the food to the foregut (Movie 3)^41^. Calcium imaging with pan-neuronal Tg(*HuC:H2B-GCaMP6s*) at 7-8 dpf, shows that posterior mXns but not anterior mXns respond to this stimulus (Movie 3, Fig. 7C, D, I). We did observe activity in neurons outside of the isl1:Kaede-expressing vagus motor nucleus, likely reflecting the complexity of the swallowing response (Fig. 7D)^42^.

**Figure 7.**
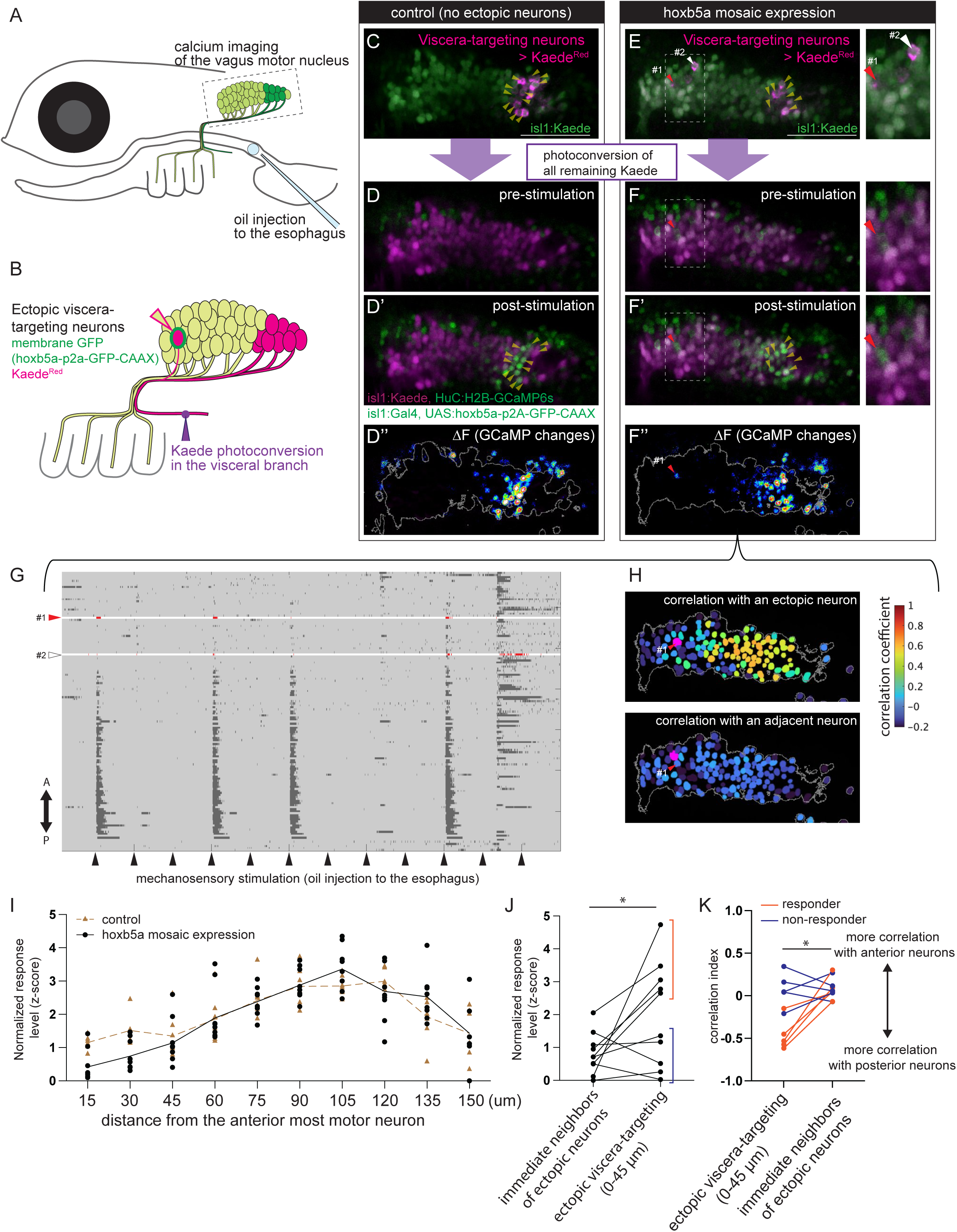
Viscera-innervating neurons can integrate into a visceral circuit independent of their position in the motor nucleus. **(A)** Schematic of esophagus stretch method for generating a specific response in posterior mXns. **(B)** Schematic of approach for generating ectopic viscera-innervating neurons by sparse expression of *hoxb5a*-GFP (green). Ectopic neurons innervating the viscera (magenta) are identified by photoconversion of the visceral branch (purple arrowhead). **(C, E)** Viscera-targeting mXns (magenta; indicated by yellow arrowheads) identified by photoconversion in control (**C**) and in larvae where hoxb5a-GFP is sparsely expressed (**E**). #1 and #2 in **E** indicate ectopic viscera-targeting mXns identified by the expression of membrane GFP and Kaede^Red^. **(D, F)** GCaMP6s signals (green) in mXns (magenta) before (**D, F**) and after (**D’, F’**) esophagus stretch in larvae shown in **C** (for **D**) and **E** (for **F**) respectively. Changes in GCaMP6s (ΔF) are shown in heat-color in **D”, F”**. Calcium increase was detected in the ectopic mXn #1 (red arrowhead). **(G)** Raster traces for mXns in the larva in **F**. Neurons aligned from top to bottom corresponding to their anterior-to-posterior position in the nucleus. Raster plots for ectopic neurons #1 and #2 are highlighted in red. **(H) Top:** computational analysis of correlation between the ectopic viscera-innervating mXn (#1) and all other mXns in the same focal plane. **Bottom:** correlation between a mXn adjacent to #1 and all other mXns. All the time frames acquired from calcium imaging were used for the correlation analysis. Heat map ranges from anti-correlated (blue) to fully correlated (red). **(I)** Normalized dF/F0 (z-score) in mXns in each of 15-µm bins along the A-P extent of the vagus motor nucleus in control (brown dotted line; n=8 larvae) and in larvae with ectopic viscera-targeting mXns (black; n=8 larvae) at 7 dpf. **(J)** Normalized dF/F0 (z-score) of individual ectopic mXns (n=10 neurons) is shown compared to the average of this value from all immediate neighboring neurons (within 8 µm). Five ectopic neurons that showed high calcium responses were categorized as responders (red bracket, including neuron #1 shown in **E**) and the others as non-responders (blue; including neuron #2 shown in **E**). **(K)** Correlation index (see Methods) was quantified for 10 ectopic neurons and their immediate neighbors analyzed in **J**. Colors indicate two groups of ectopic neurons defined in **J**, while the statistical comparison was done altogether (without separating to the two groups). Statistics in **J**, **K**: paired nonparametric t-test (Wilcoxon matched-pairs signed rank test). scale bar: 50 µm.

This selective posterior mXn activation caused by esophagus stretch provided us the opportunity to study the contribution of neural positioning to circuit wiring amongst viscera-targeting mXns. We generated ectopic viscera-targeting mXns through the sparse overexpression of *hoxb5a*, which causes anterior mXns to project in the visceral branch^8^ (Fig.7B, E). To identify these ectopic viscera-targeting mXns, we photoconverted the visceral branch of Tg(*isl1:Kaede*; *isl1:Gal4*; *UAS:hoxb5a-p2A-GFP-CAAX*; *HuC:H2B-GCaMP6s*) larvae and looked for membrane-GFP-expressing mXns in the anterior part of the motor nucleus that were retrogradely labeled with Kaede^RED^ (Fig. 7E, F). We particularly focused on the ectopic viscera-targeting mXns in the anterior 45 µm of the nucleus, where pharynx-innervating mXns reside. After identifying ectopic mXns, we photoconverted the entire motor nucleus to eliminate remaining Kaede^GREEN^ fluorescence and performed GCaMP6s imaging while injecting oil into the esophagus.

In 7-8 dpf larvae, we found that oil injections frequently activated ectopic viscera-targeting mXns in the anterior vagus motor nucleus (Fig. 7F (#1), G, J). Given the sporadic induction of motor activity following the oil injection, we analyzed the temporal correlation in neural activity between the ectopic viscera-targeting mXns and other mXns. We found that the activity pattern of ectopic anterior mXns is more correlated with posterior mXns than with anterior mXns that surround them (Fig.7H, K). Our results support that, as for ectopic pharynx-innervating mXns, the activity of ectopic viscera-targeting mXns corresponds with their axonal innervation pattern, not their position in the motor nucleus.

Taken together, our results show that both pharynx- and viscera-targeting mXns can be incorporated into correct circuits independent of their positions on the topographic map. While the limitations of the esophageal stretch experimental approach prevent us from asking whether this is an experience-dependent process involving dendritic re-targeting for viscera-targeting mXns like it is for pharynx-innervating mXns, our findings suggest that a position-independent strategy is a general principle across diverse vagus motor neuron target groups.

## DISCUSSION

This study unveiled a striking ability of vagus motor neurons to develop specific input connectivity that is appropriate for their innervation targets, independent of their position within the motor nucleus. Although vagus motor neurons that innervate different targets are spatially intermingled in the brain of larval zebrafish, they produce stereotyped peripheral outputs that differently regulate these different targets. We demonstrate remarkable resilience of input connectivity to manipulations that alter motor neuron positions and identified a single-neuron-level adaptive dendritic extension as one plausible mechanism. We additionally provide evidence that specific input connectivity is in part established by neurotransmission-mediated feedback from the periphery, supporting experience-dependent regulation. This study has established a tractable zebrafish model for investigating mechanistic insights into vagus motor wiring.

### Position-dependent and position-independent strategies in motor wiring

Our findings in the vagus system contrast with well-studied mammalian spinal motor circuits in which motor neurons innervating different muscles are generally separated into spatially discrete motor pools^4^. These receive monosynaptic input from proprioceptive sensory neurons that innervate the same muscle based on the position of the motor pool in the spinal cord^43,44^. Genetic manipulations that change the motor pool organization result in mis-wiring and consequent motor behavior defects^7^, demonstrating an indispensable role of spatial organization in specific motor wiring. We previously showed that zebrafish vagus motor neurons have a topographic relationship with their peripheral targets and hypothesized that this relationship serves to position motor neurons optimally to receive sensory input from the same body regions^8,21^. However, the lack of spatially discrete motor pools within the vagus motor nucleus predicted that a simple positional mechanism is insufficient to generate such precise connectivity. Our identification of the resilience of vagus motor neurons to topographic defects, which we have described both at the population level (Fig. 4A) and at the single-cell level (Fig. 4E), provides direct evidence of a position-independent strategy in motor wiring.

If vagus motor neurons can connect to appropriate upstream circuits independent of position in the motor nucleus, then what is the role of the topography that is established in the embryo^8,21^? Our data favor that this topographic organization instructs the coarse functional segregation among pharyngeal axonal branches that is already detectable in 4 dpf larvae (Fig. 2L). We suspect that this coarse topographic organization among pharynx-innervating neurons enables broad motor nucleus activities at 4 dpf (Fig. 3F) to result in spatially biased peripheral control. Additionally, we note that there is a limit to the distance over which topographic defects can be corrected, given that only about a half of mis-positioned neurons show successful circuit integration (non-responders in Fig7J, K), or generate adaptive dendritic projections (Fig. 5E). Thus, the position-independent strategy cannot guarantee accurate connectivity in ectopic motor neurons far away from their normal topographic range, indicating the role of coarse topographic organization as an important starting point. We propose that the position-independent strategy serves to refine the connectivity of adjacent neurons belonging to different target groups. Finally, we note that we were unable to test whether mispositioned neurons also respond to stimuli that activate neighboring neurons, a bivalent response that would be predicted to be maladaptive.

Since mammalian vagus motor neurons innervating different targets are also highly intermingled^10,11,19,31,45^, we speculate that in mammals too, vagus motor neurons may adjust their connections to presynaptic neurons to fire differently from adjacent neurons in order to generate target-specific motor responses. Furthermore, since the lack of motor pool clustering is common in other cranial motor neurons, position-independent connectivity adjustment may be a general principle in brain motor wiring. In fact, ectopic facial motor neurons can successfully generate gross motor outputs in zebrafish, though with some abnormality, when the entire facial motor population is collectively shifted with preserved topographic organization^9,46^.

### Experience-dependent connectivity refinement

Topographic organization of cell bodies is similarly coarse or absent in some sensory neurons, including mouse vagus sensory neurons^47,48^; however, specific connectivity is generally established by topographic sorting of the central axons and synapse-level adjustment. For instance, cell bodies of mouse trigeminal sensory neurons innervating neighboring whiskers are partly intermingled in the trigeminal ganglion^49^. But their central axons in the brainstem are segregated into distinct domains called barrelettes, where they synapse on different postsynaptic neurons^38^. While molecular differences play a major role in topographically sorting the central axons of whisker-innervating sensory neurons in prenatal mice, sensory experience from the periphery refines the topographic organization and instructs point-to-point synaptic connections postnatally^50,51^. Sensory experience gives correlated activity to sensory neurons innervating the same whisker and directs their central axons to converge into the same circuit through the Hebbian mechanism, enabling “neurons that fire together to wire together”^52,53^. In contrast, sensory neurons innervating different whiskers show less correlated activity and are consequently segregated into different circuits.

It is difficult to imagine how a Hebbian mechanism could refine vagus motor wiring, given that sensory input cannot alter motor axon targeting. In many primary motor neurons, including spinal and vagus motor neurons, axons travel a long distance to innervate peripheral targets early in development and cannot change them when later-developing somatodendritic connections begin generating input-induced neural activity^7,8,35,54–56^. Thus for motor circuit refinement to happen, the somatodendritic connections need to be adjusted so that motor neurons that are already wired together to common body parts fire together to generate accurate motor control. To our knowledge, such an experience-dependent refinement in *input* connectivity, in contrast to the Hebbian experience-dependent refinement of *output* connectivity has not previously been described.

We provide three lines of evidence that input connectivity of vagus motor neurons is refined in an experience-dependent manner. First, vagus motor neurons are progressively more fine-tuned to designated sensory signals after the onset of sensory-evoked activity. Second, vagus motor neurons that are made ineffective in producing motor outputs by neurotransmission suppression reduce their responsiveness to sensory signals. Third and most importantly, vagus motor neurons increase activity levels when neurotransmission is suppressed from other vagus motor neurons co-innervating the same target. This compensatory change in intact neurons strongly suggests that motor neurons that produce a relatively effective response to upstream signals are “rewarded” by a periphery-to-brain feedback signal that stabilizes input connectivity at the single neuron level. This functional compensation represents the remarkable capacity of experience-dependent refinement to help generate normal motor outputs.

The nature of this feedback signal is unknown. We predict that this signal originates from postsynaptic targets, carries information about peripheral responses to localized sensory cues, and then adjusts motor connections with upstream neurons by pruning inappropriate synapses and strengthening appropriate ones. This may be mediated by the retrograde transport of signaling molecules inside motor neurons, such as pMad transcription factor known to be activated by BMP signaling from the postsynaptic muscles in the *Drosophila* NMJ to strengthen motor synapses^57,58^. Alternatively, it could be mediated by sensory feedback, as is used in the vestibulo-occular reflex circuit to globally modify the activity level of eye-rotating motor populations^59^. How sensory feedback can modify single-neuron-level connections remains an open question, however the powerful genetic and transplantation approaches for single-cell-level manipulation in zebrafish will provide a unique system to address the nature of this non-Hebbian experience-dependent refinement.

We identified that topographically mispositioned neurons extend dendrites into the territory occupied by dendrites of motor neurons innervating the same target, which we termed “adaptive dendritic extensions” and provided evidence that the formation of adaptive dendritic extensions also depends on periphery-to-brain feedback. It is plausible that dendritic projections of vagus motor neurons are guided by feedback-mediated synaptic stabilization whereby developing motor dendrites randomly extend filopodia long distances, and nascent connections that produce relatively effective motor outputs are more stabilized than others. Such a mechanism was previously proposed based on live imaging of zebrafish tectal neurons^60^. This step could guide the direction of dendritic projections and eventually create dendritic architecture that topographically echoes the already-fixed axonal projection patterns of vagus motor neurons.

Even in mammalian spinal motor circuits mentioned above, motor neurons innervating synergistic muscles are spatially intermingled, and their precise synaptic connectivity with upstream sensory axons requires neural activities^61^. We speculate that a periphery-to-CNS feedback mechanism we identified in the vagus nerve is also used in spinal motor circuits for fine-scale connection specificity.

### Vagus sensory-motor circuits across vertebrates

In this study, we investigated input connectivity of vagus motor neurons by analyzing their activity responses to sensory signals. The anatomical correlates of these responses are poorly understood in any system, including in the zebrafish where the direct presynaptic partners of vagus motor neurons are unidentified. Reportedly, many reflexes that involve both sensory and motor components of the vagus are controlled by a 3-neuron circuit (i.e., sensory neuron, interneuron, motor neuron)^15,28,62,63^. Both in carp and catfish, such simple 3-neuron reflex arcs connect vagus sensory and motor neurons innervating the same pharyngeal locations to generate the oropharyngeal reflex that sorts and expels inedible items from the pharynx^28,63^. The stereotypical pharyngeal reactions against pharyngeal noxious cues we observe in zebrafish larvae may likewise be mediated by a 3-neuron arc generated from vagus sensory and motor neurons innervating the pharynx. If this is indeed the case, identifying presynaptic neurons of vagus motor neurons would reveal all components of the sensory-motor circuit and allow us to investigate the anatomical basis of specific motor wiring. Future studies will determine how single interneurons in the brainstem identify and bridge functionally appropriate pairs of sensory and motor neurons of the vagus for functional reflex circuit wiring. This question is important because the 3-neuron structure is widely conserved even in mammals for some vagus-mediated reflexes, which include the gastric and pancreatic reflexes^15,62^. It remains poorly explored in any animal how postsynaptic neurons of vagus sensory neurons in the nucleus tractus solitarii (NTS) synapse on specific vagus motor neurons for independent body controls, though remarkable functional topography of NTS neurons was recently reported^64^. The zebrafish system we established in this study is particularly suitable for addressing this question.

While the simple 3-neuron arc is partly conserved across vertebrates, vagus reflex circuits have modified networks to optimize peripheral body reactions as vertebrates evolved internal body structures and functions. In mouse, pharyngeal muscle activities are reflexively induced by sensory cues in the upper airway for airway protection^65^. Additionally, mammals respond to inflammation-driven noxious signals in the pharynx, not with simple pharyngal muscle contractions but with the complex cough reflex which is regulated by compound vagus-involved networks in the medulla called the cough center^66^. We hypothesize that acquisition of optimized vagus networks during vertebrate evolution has been accelerated by the plasticity we illustrated in this study, which can modify motor connectivity in the absence of prior genetic adjustment in molecular neuron-neuron recognition.

## ACKNOWLEDGMENTS

We thank Drs. Ajay Dhaka, Herwig Baier, and Koichi Kawakami and National Bioresource Project of Japan for transgenic zebrafish lines. We thank Dr. Shin-Ichi Higashijima for DNA plasmids. We thank Dr. Penny Lam for assistance with optovin-mediated noxious stimulation in larval zebrafish. We thank Drs. Akira Muto and Koichi Kawakami for initial assistance with hindbrain calcium imaging. We thank Drs. Bing Ye, David Schoppik, Jeremy Dasen, Michael Granato and Virginia MS Ruetten for critical feedback on an earlier version of the manuscript. We thank all the members of the Moens lab for discussion, Jason Stonick for technical help, and Olivia Fitzgerald for zebrafish care. This work was supported by NIH grants R01 (NS109425) and R21 (NS124191) to C.B.M., F32 (HD096860) to A.J.I, and an AHA postdoctoral fellowship (19POST34380845), a JSPS overseas research fellowship (202060651), and NIG-JOINT (National Institute of Genetics: 96I2019) to T.K.

## Author Contribution

T.K. and C.B.M. conceived the project. T.K. performed the experiments. J.B-W. wrote scripts for automated analysis. T.K. and A.J.I. generated transgenic zebrafish lines. T.K. and C.B.M. wrote the paper.

## METHODS AND MATERIALS

### Experimental Model and Subject Details

#### Zebrafish lines

*Danio rerio* animals were raised at the Fred Hutchinson Cancer Center in accordance with institutional animal care and use committee (IACUC)-approved protocols. All experiments were carried out in accordance with IACUC standards. Fish were bred and maintained according to standard protocols. All experimental stages are noted in the figures and text. Sex is not a relevant biological variable in our experiments, which are carried out before sex is determined in zebrafish. Larval zebrafish were fed starting at 5 dpf. Transgenic lines used in this study include: *Tg(isl1:Kaede)^ch103^* ^8^ ; *Tg(isl1:Gal4)^fh452^* ^67^; *Tg(isl1:mRFP)^fh1^* ^68^ ; *Tg(isl1:EGFPCAAX)^fh474^* ^8^; *Tg(p2rx3b:GFP)^sl1^* ^69^ *TgBAC(chata:GAL4-VP16)^mpn202^* ^70^; *Tg(UAS-E1B:Kaede)^s1999t^* ^71^; Tg(*UAS:GCaMP6s^13A^)* ^72^; *Tg(UAS:BoTxBLC-GFP^icm21^)* ^36^; *HuC:H2B-GCaMP6s* ^73^. Lines generated for this study include *Tg(isl1:H2B-GCaMP6s)^fh579,^ Tg(UAS:hoxb5a-p2a-GFP-CAAX)^fh567^, Tg(UAS:DN-RAR-p2a-GFP-CAAX)^fh575^, Tcf21^Gal4(fh557)^,* and *Trpa1b^Gal4(fh577)^*.

### Method Details

#### Generation of transgenic lines

50 pg of the Tol2 plasmid *UAS:hoxb5a-p2a-GFP-CAAX or UAS:DN-RAR-p2a-GFP-CAAX,* which was generated previously^8,21^, was injected into *Tg(isl1:Gal4)* embryos at the one-cell stage together with 50 pg of Tol2 transposase mRNA. The Tol2 plasmid *isl1:H2B-GCaMP6s* was generated in this study. The *H2B-GCaMP6s* sequence was obtained from the Addgene plasmid *Tol2-elav3-H2B-GCaMP6s* (#59530), and the *isl1* CREST1 enhancer is from a previous study^30^. 50 pg of the Tol2 plasmid *isl1:H2B-GCaMP6s* was injected into wild type embryos at the one-cell stage together with 50 pg of Tol2 transposase mRNA. Injected embryos were screened for GFP-CAAX or GCaMP6s expression and grown to adulthood. Stable lines were then identified in the F1 generation.

*Tcf21^Gal4^, Trpa1b^Gal4^,* were generated using the CRISPR/Cas9-mediated knock-in strategy outlined in^74^. The donor plasmids GBait-hs-Gal4 were provided by the Higashijima lab.

#### Kaede photoconversion

Kaede photoconversion to visualize target groups of vagus motor neurons was described previously ^8,21^. In brief, larvae were anesthetized with 200 mg/l 3-aminobenzoic acid ethyl ester (MS-222) (Sigma-Aldrich, A5040) and embedded laterally in 1.2% low-melting-point (LMP) agarose (Invitrogen, 16520). A region of interest (ROI) drawn over an individual branch was photoconverted by 100 iterations of 405 nm laser at 8% power on a Zeiss LSM 700 inverted confocal microscope. Kaede^Red^ signal in the vagus motor nucleus was imaged at least 1 hr after the photoconversion on a Zeiss LSM 700 or Leica Stellaris5 confocal microscope.

For sequential photoconversion in Fig 1C and FigS1, the cycle of photoconversion and image acquisition was repeated for different branches in the same animal, with at least 1 hr between each photoconversion and imaging. To determine Kaede^Red^ signals generated by the latest photoconversion but not by the prior photoconversions, one preceding Kaede^Red^ image was subtracted from the latest Kaede^red^ image. To subtract an image, the Image Calculator function of Fiji was used after Kaede^Red^ images from different time points were aligned by Descriptor-based registration function of Fiji.

#### Calcium imaging

Calcium imaging was done on Zeiss LSM700 confocal microscope with single z-plain acquisition at 4 Hz. For focal noxious stimulation, larval zebrafish were incubated with 10 µM optovin dissolved in 0.2% DMSO for at least 2 hours prior to calcium imaging. Larvae were then embedded laterally in 1.2% LMP agarose without tricaine treatment. To focally activate optovin, 405 nm confocal laser at 6% power was given for ∼2 seconds to a region of interest drawn over the anterior pharynx, posterior pharynx, or the esophagus. Photostimulation was repeated within the same animal at 25-40 second intervals (100 frames for nucleus imaging and 150 frames for axonal imaging). For AITC-mediated noxious stimulation, 10 mM AITC dissolved in 1% DMSO was bath applied onto agarose-mounted animals during calcium imaging.

To provide mechanosensory stimulation to the larval esophagus, a glass needle pulled from a Wiretrol 10 µl capillary micropipette (Drummond Scientific, 21-175B) was filled with mineral oil (Sigma) and inserted into the esophagus through the stomach, in laterally agarose-embedded larval zebrafish without tricaine treatment while on the confocal microscope stage. During calcium imaging from the motor nucleus, drops of mineral oil (approximately 30 µm in diameter) were delivered inside the esophagus every 25 seconds (every 100 frames). To identify the viscera-targeting neurons in animals that carry Tg*(isl1:Kaede)*, Kaede was photoconverted in the visceral branch, and Kaede^Red^ was imaged at least 1 hour after the photoconversion. All remaining Kaede^Green^ signals were photoconverted to Kaede^Red^ prior to GCaMP imaging.

#### Vagus neuron transplantation

Transplantation was performed as described previously^8,35^ on 2 dpf embryos in Fig. 4 and 1 dpf embryos for Fig. 6G-J. Donor embryos carrying Tg(*isl1:mRFP*) and Tg(*isl1:H2B-GCaMP6s*) (nuclear GCaMP6s) were used in Fig. 4E-O and Fig. 6G-J for nuclear GCaMP imaging, whereas donor embryos carrying Tg(*isl1:mRFP*), Tg(*chata:Gal4*) and Tg(*UAS:GCaMP6s*) (cytoplasmic GCaMP6s) were used in Fig. 4A-D for axonal GCaMP imaging. Host embryos carrying Tg(*isl1:Gal4*, *UAS-BoTxBLC-GFP*) were used in Fig. 4E-O and Fig. 6G-J, whereas Tg(*chata:Gal4*); Tg(*UAS-BoTxBLC-GFP*) was used in Fig. 4A-D for axonal GCaMP imaging because isl1:Gal4-driven GFP expression in the heart overlaps with the GCaMP signals in motor axons. For transplantation, donor and host embryos were both anesthetized with 200 mg/l MS-222 and laterally embedded in 1.2 % LMP agarose. A Wiretrol 10 µl capillary micropipette (Drummond Scientific, 21-175B) was pulled and the tip broken to make a 10 µm opening. The needle was inserted from the dorsal side into the hindbrain at a 45° angle to the body axis at the level of the vagus nucleus. RFP-expressing vagus neurons were collected from the particular topographic position of the donor nucleus stated in the text. The needle was then similarly inserted into the vagus nucleus of host embryos, and ∼10 RFP-neurons were placed into a designated position of the host nucleus. Following transplantation, larvae were unmounted from agarose and placed into normal Ringer’s solution+1% Pen Strep (Gibco, 15140-122) until imaging at 10-11 dpf. Because a majority of neurons transplanted after 40 hpf disappeared by 10 dpf or showed no calcium transients at all, neurons that showed at least one calcium transient were used for quantification.

#### Mosaic transgenic labeling for dendritic imaging

UAS:DN-RAR-p2A-GFP-CAAX (Fig. 5) and UAS:hoxb5a-p2a-GFP-CAAX (Fig. 7) were sparsely expressed in vagus motor neurons using a Tg(*isl1:Gal4*) driver in highly variegated stable transgenic lines. Since our Tg(*UAS:GFP-CAAX*) stable line is not variegating to the same degree, we generated sparsely GFP-CAAX expressing control mXns (Fig. 5A, B) by plasmid injection as described previously^8^. In brief, one-cell-stage Tg(*isl1:Gal4*) embryos were injected with 50 pg of *Tol2* mRNA, combined with 25 pg of UAS:GFP-CAAX plasmid^8^. Injected animals were screened for sparse GFP expression at 3 dpf and raised to 7-8 dpf for confocal imaging.

### Quantification and Statistical Analysis

#### Quantification of calcium responses in the motor nucleus

Calcium imaging data of the vagus motor nucleus were first motion-corrected by CaImAn^75^, followed by automated cellular segmentation by the machine-learning-based method Cellpose^76^. Significant calcium transients in individual regions of interest (ROIs) were then detected by the MatLab-based toolbox developed previously^77^ where (1) the noise in the fluorescent signal was estimated from the standard deviation of the baseline fluorescent variations and (2) *Dynamic Threshold Method* was used to estimate the significant fluorescent transients. Mean dF/F0 from the 20 frames following noxious stimulation were shown in Fig. 3D, I and used for pair-wise correlation analysis in Fig. 3E. In esophageal stretch experiments, larvae that show stretch-induced responses over 3 times within 2000 time frames were used for data analysis. Because ROIs for hoxb5a-positive neurons in Fig. 7I, J, contain GFP-CAAX-derived green fluorescent signals, to compare changes in GCaMP fluorescence in GFP-CAAX-positive (hoxb5a-positive) neurons and GFP-CAAX-negative neighbors, dF/F0 values are normalized into the Z-score and averaged for all time frames when >50% posterior neurons show significant transients. Temporal correlation in significant transients in Fig. 7H, K was analyzed for correlation coefficient between a single ROI and all other ROIs (vagus motor neurons in the same imaging plane). The correlation coefficient that single anterior neurons (either ectopic or not) show with all other anterior neurons (0∼45 µm, where pharynx-innervating neurons reside) and posterior neurons (75 µm∼) were each averaged for Corr^Ant^ and Corr^Post^, respectively. Correlation index in Fig 7K was calculated as (Corr^Ant^ – Corr^Post^)/(Corr^Ant^+Corr^Post^), where Wilcoxon matched-pairs singled rank test was used.

#### Quantification of calcium responses in pharyngeal arches

GCaMP signals were measured from ROIs drawn manually over the pharyngeal arches PA4, 5, 6 as indicated in Fig. 2B. Mean dF/F0 values from the 10 frames following noxious stimulation were plotted in Fig 2D, H, K, where paired t-test was used. Response tuning index was calculated for individual ROIs as [(dF^A^/F0)– (dF^P^/F0)] / [(dF^A^/F0) + (dF^P^/F0)], where dF^A^/F0 and dF^P^/F0 indicate mean dF/F0 following noxious stimulation to the anterior and posterior pharynx, respectively. Paired nonparametric test (Wilcoxon matched-pairs signed rank test) was used.

#### Quantification of calcium responses in vagus sensory axons

To compare the axonal activity in vagus sensory neurons elicited by optovin stimulation to different body locations, cumulative density distribution of dF/F0 was measured. To calculate dF/F0 for each pixel, the 20 frames prior to noxious stimulation were averaged into a projection image, “pre-image”, and the 20 frames after stimulation were similarly averaged into a projection image, “post-image”. Image Calculator function of Fiji was used to subtract the pre-image from the post-image for dF, which was then divided by the pre-image, generating a 2D image with dF/F0 in each pixel. Cumulative density distribution was then quantified for the anterior-posterior extent of the axon terminals as described previously^35^. The half-maximum value of the cumulative density distribution was plotted in Fig. S2I and statistically analyzed by t-test.

#### Quantification of adaptive dendritic extensions

In Fig. 4O, the proportion of the dendritic area anterior to the cell body as a fraction of the total dendritic area was plotted and compared by t-test. The dendritic area was measured as a pixel count after automated thresholding of dendrite images (e.g., magenta in Fig. 4H, L). In Fig. 5E, GFP-labeled posterior mXns whose dendrites extend beyond the posterior 40% of the dendritic neuropil were defined as “neurons with anterior dendritic projections”, and chi-square test was used to compare across conditions. In Fig. 5F, the A-P distance along the longitudinal dendritic neuropil between the anterior dendritic limit of GFP-labeled motor neurons and the position of the cell body was measured and compared across conditions through Mann-Whitney test.

## REFERENCES

1 Cang, J. & Feldheim, D. A. Developmental mechanisms of topographic map formation and alignment. Annu Rev Neurosci 36, 51–77, doi:10.1146/annurev-neuro-062012-170341 (2013).

2 Schreiner, C. E. & Winer, J. A. Auditory cortex mapmaking: principles, projections, and plasticity. Neuron 56, 356–365, doi:10.1016/j.neuron.2007.10.013 (2007).

3 Erzurumlu, R. S., Murakami, Y. & Rijli, F. M. Mapping the face in the somatosensory brainstem. Nat Rev Neurosci 11, 252–263, doi:10.1038/nrn2804 (2010).

4 Levine, A. J., Lewallen, K. A. & Pfaff, S. L. Spatial organization of cortical and spinal neurons controlling motor behavior. Curr Opin Neurobiol 22, 812–821, doi:10.1016/j.conb.2012.07.002 (2012).

5 Balaskas, N., Abbott, L. F., Jessell, T. M. & Ng, D. Positional Strategies for Connection Specificity and Synaptic Organization in Spinal Sensory-Motor Circuits. Neuron 102, 1143–1156 e1144, doi:10.1016/j.neuron.2019.04.008 (2019).

6 Tripodi, M. & Arber, S. Regulation of motor circuit assembly by spatial and temporal mechanisms. Curr Opin Neurobiol 22, 615–623, doi:10.1016/j.conb.2012.02.011 (2012).

7 Surmeli, G., Akay, T., Ippolito, G. C., Tucker, P. W. & Jessell, T. M. Patterns of spinal sensory-motor connectivity prescribed by a dorsoventral positional template. Cell 147, 653–665, doi:10.1016/j.cell.2011.10.012 (2011).

8 Barsh, G. R., Isabella, A. J. & Moens, C. B. Vagus Motor Neuron Topographic Map Determined by Parallel Mechanisms of hox5 Expression and Time of Axon Initiation. Curr Biol 27, 3812–3825 e3813, doi:10.1016/j.cub.2017.11.022 (2017).

9 McArthur, K. L. & Fetcho, J. R. Key Features of Structural and Functional Organization of Zebrafish Facial Motor Neurons Are Resilient to Disruption of Neuronal Migration. Curr Biol 27, 1746–1756 e1745, doi:10.1016/j.cub.2017.05.033 (2017).

10 Hopkins, D. A., Bieger, D., deVente, J. & Steinbusch, W. M. Vagal efferent projections: viscerotopy, neurochemistry and effects of vagotomy. Prog Brain Res 107, 79–96, doi:10.1016/s0079-6123(08)61859-2 (1996).

11 Kitamura, S., Ogata, K., Nishiguchi, T., Nagase, Y. & Shigenaga, Y. Location of the motoneurons supplying the rabbit pharyngeal constrictor muscles and the peripheral course of their axons: a study using the retrograde HRP or fluorescent labeling technique. Anat Rec 229, 399–406, doi:10.1002/ar.1092290312 (1991).

12 Tenney, A. P. et al. Etv1 Controls the Establishment of Non-overlapping Motor Innervation of Neighboring Facial Muscles during Development. Cell Rep 29, 437–452 e434, doi:10.1016/j.celrep.2019.08.078 (2019).

13 Neuhuber, W. L. & Berthoud, H. R. Functional anatomy of the vagus system - Emphasis on the somato-visceral interface. Auton Neurosci 236, 102887, doi:10.1016/j.autneu.2021.102887 (2021).

14 Prescott, S. L. & Liberles, S. D. Internal senses of the vagus nerve. Neuron 110, 579–599, doi:10.1016/j.neuron.2021.12.020 (2022).

15 Mussa, B. M. & Verberne, A. J. The dorsal motor nucleus of the vagus and regulation of pancreatic secretory function. Exp Physiol 98, 25–37, doi:10.1113/expphysiol.2012.066472 (2013).

16 Coverdell, T. C., Abbott, S. B. G. & Campbell, J. N. Molecular cell types as functional units of the efferent vagus nerve. Semin Cell Dev Biol, doi:10.1016/j.semcdb.2023.07.007 (2023).

17 Yuan, H. & Silberstein, S. D. Vagus Nerve and Vagus Nerve Stimulation, a Comprehensive Review: Part I. Headache 56, 71–78, doi:10.1111/head.12647 (2016).

18 Isabella, A. J. & Moens, C. B. Development and regeneration of the vagus nerve. Semin Cell Dev Biol, doi:10.1016/j.semcdb.2023.07.008 (2023).

19 Bieger, D. & Hopkins, D. A. Viscerotopic representation of the upper alimentary tract in the medulla oblongata in the rat: the nucleus ambiguus. J Comp Neurol 262, 546–562, doi:10.1002/cne.902620408 (1987).

20 Fox, E. A. & Powley, T. L. Longitudinal columnar organization within the dorsal motor nucleus represents separate branches of the abdominal vagus. Brain Res 341, 269–282, doi:10.1016/0006-8993(85)91066-2 (1985).

21 Isabella, A. J., Barsh, G. R., Stonick, J. A., Dubrulle, J. & Moens, C. B. Retinoic Acid Organizes the Zebrafish Vagus Motor Topographic Map via Spatiotemporal Coordination of Hgf/Met Signaling. Dev Cell 53, 344–357 e345, doi:10.1016/j.devcel.2020.03.017 (2020).

22 Veerakumar, A., Yung, A. R., Liu, Y. & Krasnow, M. A. Molecularly defined circuits for cardiovascular and cardiopulmonary control. Nature 606, 739–746, doi:10.1038/s41586-022-04760-8 (2022).

23 Kirshner, H. S. Causes of neurogenic dysphagia. Dysphagia 3, 184–188, doi:10.1007/BF02407221 (1989).

24 Santoso, L. F., Kim, D. Y. & Paydarfar, D. Sensory dysphagia: A case series and proposed classification of an under recognized swallowing disorder. Head Neck 41, E71–E78, doi:10.1002/hed.25588 (2019).

25 Cunningham, E. T., Jr., Ravich, W. J., Jones, B. & Donner, M. W. Vagal reflexes referred from the upper aerodigestive tract: an infrequently recognized cause of common cardiorespiratory responses. Ann Intern Med 116, 575–582, doi:10.7326/0003-4819-116-7-575 (1992).

26 Cutsforth-Gregory, J. K. & Benarroch, E. E. Nucleus of the solitary tract, medullary reflexes, and clinical implications. Neurology 88, 1187–1196, doi:10.1212/WNL.0000000000003751 (2017).

27 Herbst, J. J., Minton, S. D. & Book, L. S. Gastroesophageal reflux causing respiratory distress and apnea in newborn infants. J Pediatr 95, 763–768, doi:10.1016/s0022-3476(79)80733-7 (1979).

28 Finger, T. E. Evolution of gustatory reflex systems in the brainstems of fishes. Integr Zool 4, 53-63, doi:10.1111/j.1749-4877.2008.00135.x (2009).

29 Ikenaga, T., Ogura, T. & Finger, T. E. Vagal gustatory reflex circuits for intraoral food sorting behavior in the goldfish: cellular organization and neurotransmitters. J Comp Neurol 516, 213–225, doi:10.1002/cne.22097 (2009).

30 Higashijima, S., Hotta, Y. & Okamoto, H. Visualization of cranial motor neurons in live transgenic zebrafish expressing green fluorescent protein under the control of the islet-1 promoter/enhancer. J Neurosci 20, 206–218, doi:10.1523/JNEUROSCI.20-01-00206.2000 (2000).

31 Altschuler, S. M., Bao, X. M. & Miselis, R. R. Dendritic architecture of nucleus ambiguus motoneurons projecting to the upper alimentary tract in the rat. J Comp Neurol 309, 402–414, doi:10.1002/cne.903090309 (1991).

32 Kokel, D. et al. Photochemical activation of TRPA1 channels in neurons and animals. Nat Chem Biol 9, 257–263, doi:10.1038/nchembio.1183 (2013).

33 Lee, G. H., Chang, M. Y., Hsu, C. H. & Chen, Y. H. Essential roles of basic helix-loop-helix transcription factors, Capsulin and Musculin, during craniofacial myogenesis of zebrafish. Cell Mol Life Sci 68, 4065–4078, doi:10.1007/s00018-011-0637-2 (2011).

34 Balaskas, N., Ng, D. & Zampieri, N. The Positional Logic of Sensory-Motor Reflex Circuit Assembly. Neuroscience 450, 142–150, doi:10.1016/j.neuroscience.2020.04.038 (2020).

35 Isabella, A. J., Stonick, J. A., Dubrulle, J. & Moens, C. B. Intrinsic positional memory guides target-specific axon regeneration in the zebrafish vagus nerve. Development 148, doi:10.1242/dev.199706 (2021).

36 Sternberg, J. R. et al. Optimization of a Neurotoxin to Investigate the Contribution of Excitatory Interneurons to Speed Modulation In Vivo. Curr Biol 26, 2319–2328, doi:10.1016/j.cub.2016.06.037 (2016).

37 Cline, H. T., Lau, M. & Hiramoto, M. Activity-dependent Organization of Topographic Neural Circuits. Neuroscience 508, 3–18, doi:10.1016/j.neuroscience.2022.11.032 (2023).

38 Kitazawa, T. & Rijli, F. M. Barrelette map formation in the prenatal mouse brainstem. Curr Opin Neurobiol 53, 210–219, doi:10.1016/j.conb.2018.09.008 (2018).

39 Zou, D. J. et al. Postnatal refinement of peripheral olfactory projections. Science 304, 1976–1979, doi:10.1126/science.1093468 (2004).

40 Sigrist, S. J., Reiff, D. F., Thiel, P. R., Steinert, J. R. & Schuster, C. M. Experience-dependent strengthening of Drosophila neuromuscular junctions. J Neurosci 23, 6546–6556, doi:10.1523/JNEUROSCI.23-16-06546.2003 (2003).

41 Nikaki, K., Sawada, A., Ustaoglu, A. & Sifrim, D. Neuronal Control of Esophageal Peristalsis and Its Role in Esophageal Disease. Curr Gastroenterol Rep 21, 59, doi:10.1007/s11894-019-0728-z (2019).

42 Jean, A. Brain stem control of swallowing: neuronal network and cellular mechanisms. Physiol Rev 81, 929–969, doi:10.1152/physrev.2001.81.2.929 (2001).

43 Imai, F. & Yoshida, Y. Molecular mechanisms underlying monosynaptic sensory-motor circuit development in the spinal cord. Dev Dyn 247, 581–587, doi:10.1002/dvdy.24611 (2018).

44 Shadrach, J. L., Gomez-Frittelli, J. & Kaltschmidt, J. A. Proprioception revisited: where do we stand? Curr Opin Physiol 21, 23–28, doi:10.1016/j.cophys.2021.02.003 (2021).

45 Tao, J. et al. Highly selective brain-to-gut communication via genetically defined vagus neurons. Neuron 109, 2106–2115 e2104, doi:10.1016/j.neuron.2021.05.004 (2021).

46 Asante, E. et al. Defective Neuronal Positioning Correlates With Aberrant Motor Circuit Function in Zebrafish. Front Neural Circuits 15, 690475, doi:10.3389/fncir.2021.690475 (2021).

47 Williams, E. K. et al. Sensory Neurons that Detect Stretch and Nutrients in the Digestive System. Cell 166, 209–221, doi:10.1016/j.cell.2016.05.011 (2016).

48 Chang, R. B., Strochlic, D. E., Williams, E. K., Umans, B. D. & Liberles, S. D. Vagal Sensory Neuron Subtypes that Differentially Control Breathing. Cell 161, 622–633, doi:10.1016/j.cell.2015.03.022 (2015).

49 da Silva, S. et al. Proper formation of whisker barrelettes requires periphery-derived Smad4-dependent TGF-beta signaling. Proc Natl Acad Sci U S A 108, 3395–3400, doi:10.1073/pnas.1014411108 (2011).

50 Lee, L. J., Lo, F. S. & Erzurumlu, R. S. NMDA receptor-dependent regulation of axonal and dendritic branching. J Neurosci 25, 2304–2311, doi:10.1523/JNEUROSCI.4902-04.2005 (2005).

51 Lo, F. S. & Erzurumlu, R. S. Sensory Activity-Dependent and Sensory Activity-Independent Properties of the Developing Rodent Trigeminal Principal Nucleus. Dev Neurosci 38, 163–170, doi:10.1159/000446395 (2016).

52 Hebb, D. O. The organization of behavior : a neuropsychological theory. (Wiley, 1949). 53

53. Katz, L. C. & Shatz, C. J. Synaptic activity and the construction of cortical circuits. Science 274, 1133–1138, doi:10.1126/science.274.5290.1133 (1996).

54 Bonanomi, D. & Pfaff, S. L. Motor axon pathfinding. Cold Spring Harb Perspect Biol 2, a001735, doi:10.1101/cshperspect.a001735 (2010).

55 Bonanomi, D. Axon pathfinding for locomotion. Semin Cell Dev Biol 85, 26–35, doi:10.1016/j.semcdb.2017.11.014 (2019).

56 Cox, J. A., Lamora, A., Johnson, S. L. & Voigt, M. M. Diverse mechanisms for assembly of branchiomeric nerves. Dev Biol 357, 305–317, doi:10.1016/j.ydbio.2011.06.044 (2011).

57 Berke, B., Wittnam, J., McNeill, E., Van Vactor, D. L. & Keshishian, H. Retrograde BMP signaling at the synapse: a permissive signal for synapse maturation and activity-dependent plasticity. J Neurosci 33, 17937–17950, doi:10.1523/JNEUROSCI.6075-11.2013 (2013).

58 Baines, R. A. Synaptic strengthening mediated by bone morphogenetic protein-dependent retrograde signaling in the Drosophila CNS. J Neurosci 24, 6904–6911, doi:10.1523/JNEUROSCI.1978-04.2004 (2004).

59 Ito, M. Cerebellar learning in the vestibulo-ocular reflex. Trends Cogn Sci 2, 313–321, doi:10.1016/s1364-6613(98)01222-4 (1998).

60 Niell, C. M., Meyer, M. P. & Smith, S. J. In vivo imaging of synapse formation on a growing dendritic arbor. Nat Neurosci 7, 254–260, doi:10.1038/nn1191 (2004).

61 Mendelsohn, A. I., Simon, C. M., Abbott, L. F., Mentis, G. Z. & Jessell, T. M. Activity Regulates the Incidence of Heteronymous Sensory-Motor Connections. Neuron 87, 111–123, doi:10.1016/j.neuron.2015.05.045 (2015).

62 Travagli, R. A., Hermann, G. E., Browning, K. N. & Rogers, R. C. Brainstem circuits regulating gastric function. Annu Rev Physiol 68, 279–305, doi:10.1146/annurev.physiol.68.040504.094635 (2006).

63 Finger, T. E. Sorting food from stones: the vagal taste system in Goldfish, Carassius auratus. J Comp Physiol A Neuroethol Sens Neural Behav Physiol 194, 135–143, doi:10.1007/s00359-007-0276-0 (2008).

64 Ran, C., Boettcher, J. C., Kaye, J. A., Gallori, C. E. & Liberles, S. D. A brainstem map for visceral sensations. Nature 609, 320–326, doi:10.1038/s41586-022-05139-5 (2022).

65 Prescott, S. L., Umans, B. D., Williams, E. K., Brust, R. D. & Liberles, S. D. An Airway Protection Program Revealed by Sweeping Genetic Control of Vagal Afferents. Cell 181, 574–589 e514, doi:10.1016/j.cell.2020.03.004 (2020).

66 Polverino, M. et al. Anatomy and neuro-pathophysiology of the cough reflex arc. Multidiscip Respir Med 7, 5, doi:10.1186/2049-6958-7-5 (2012).

67 Davey, C. F., Mathewson, A. W. & Moens, C. B. PCP Signaling between Migrating Neurons and their Planar-Polarized Neuroepithelial Environment Controls Filopodial Dynamics and Directional Migration. PLoS Genet 12, e1005934, doi:10.1371/journal.pgen.1005934 (2016).

68 Grant, P. K. & Moens, C. B. The neuroepithelial basement membrane serves as a boundary and a substrate for neuron migration in the zebrafish hindbrain. Neural Dev 5, 9, doi:10.1186/1749-8104-5-9 (2010).

69 Gau, P. et al. The zebrafish ortholog of TRPV1 is required for heat-induced locomotion. J Neurosci 33, 5249–5260, doi:10.1523/JNEUROSCI.5403-12.2013 (2013).

70 Forster, D. et al. Genetic targeting and anatomical registration of neuronal populations in the zebrafish brain with a new set of BAC transgenic tools. Sci Rep 7, 5230, doi:10.1038/s41598-017-04657-x (2017).

71 Scott, E. K. et al. Targeting neural circuitry in zebrafish using GAL4 enhancer trapping. Nat Methods 4, 323–326, doi:10.1038/nmeth1033 (2007).

72 Muto, A. et al. Activation of the hypothalamic feeding centre upon visual prey detection. Nat Commun 8, 15029, doi:10.1038/ncomms15029 (2017).

73 Esancy, K. et al. A zebrafish and mouse model for selective pruritus via direct activation of TRPA1. Elife 7, doi:10.7554/eLife.32036 (2018).

74 Kimura, Y., Hisano, Y., Kawahara, A. & Higashijima, S. Efficient generation of knock-in transgenic zebrafish carrying reporter/driver genes by CRISPR/Cas9-mediated genome engineering. Sci Rep 4, 6545, doi:10.1038/srep06545 (2014).

75 Giovannucci, A. et al. CaImAn an open source tool for scalable calcium imaging data analysis. Elife 8, doi:10.7554/eLife.38173 (2019).

76 Stringer, C., Wang, T., Michaelos, M. & Pachitariu, M. Cellpose: a generalist algorithm for cellular segmentation. Nat Methods 18, 100–106, doi:10.1038/s41592-020-01018-x (2021).

77 Romano, S. A. et al. An integrated calcium imaging processing toolbox for the analysis of neuronal population dynamics. PLoS Comput Biol 13, e1005526, doi:10.1371/journal.pcbi.1005526 (2017).

