## Supplemental information for "Position-independent functional refinement within the vagus motor topographic map"

### Figure S1

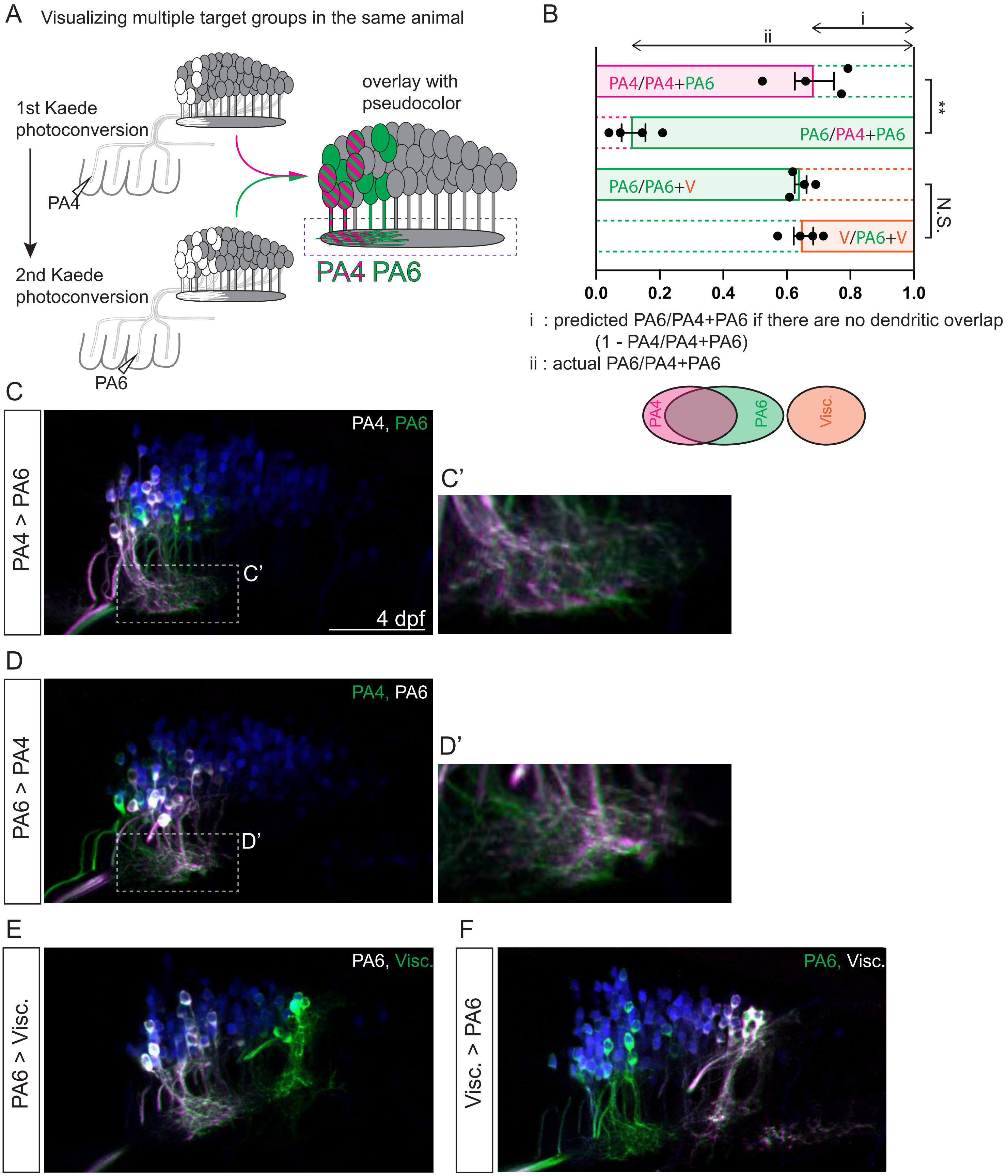
A
Visualizing multiple target groups in the same animal
1st Kaede photoconversion
PA4
2nd Kaede photoconversion
PA6
overlay with pseudocolor
PA4 PA6
B
i
ii
PA4/PA4+PA6
PA6/PA4+PA6
PA6/PA6+V
V/PA6+V
**
N.S.
0.0
0.2
0.4
0.6
0.8
1.0
i : predicted PA6/PA4+PA6 if there are no dendritic overlap
(1 - PA4/PA4+PA6)
ii : actual PA6/PA4+PA6
PA4
PA6
Visc.
C
PA4 > PA6
PA4, PA6
C'
4 dpf
C'
D
PA6 > PA4
PA4, PA6
D'
D'
E
PA6 > Visc.
PA6, Visc.
F
Visc. > PA6
PA6, Visc.

### Figure S2

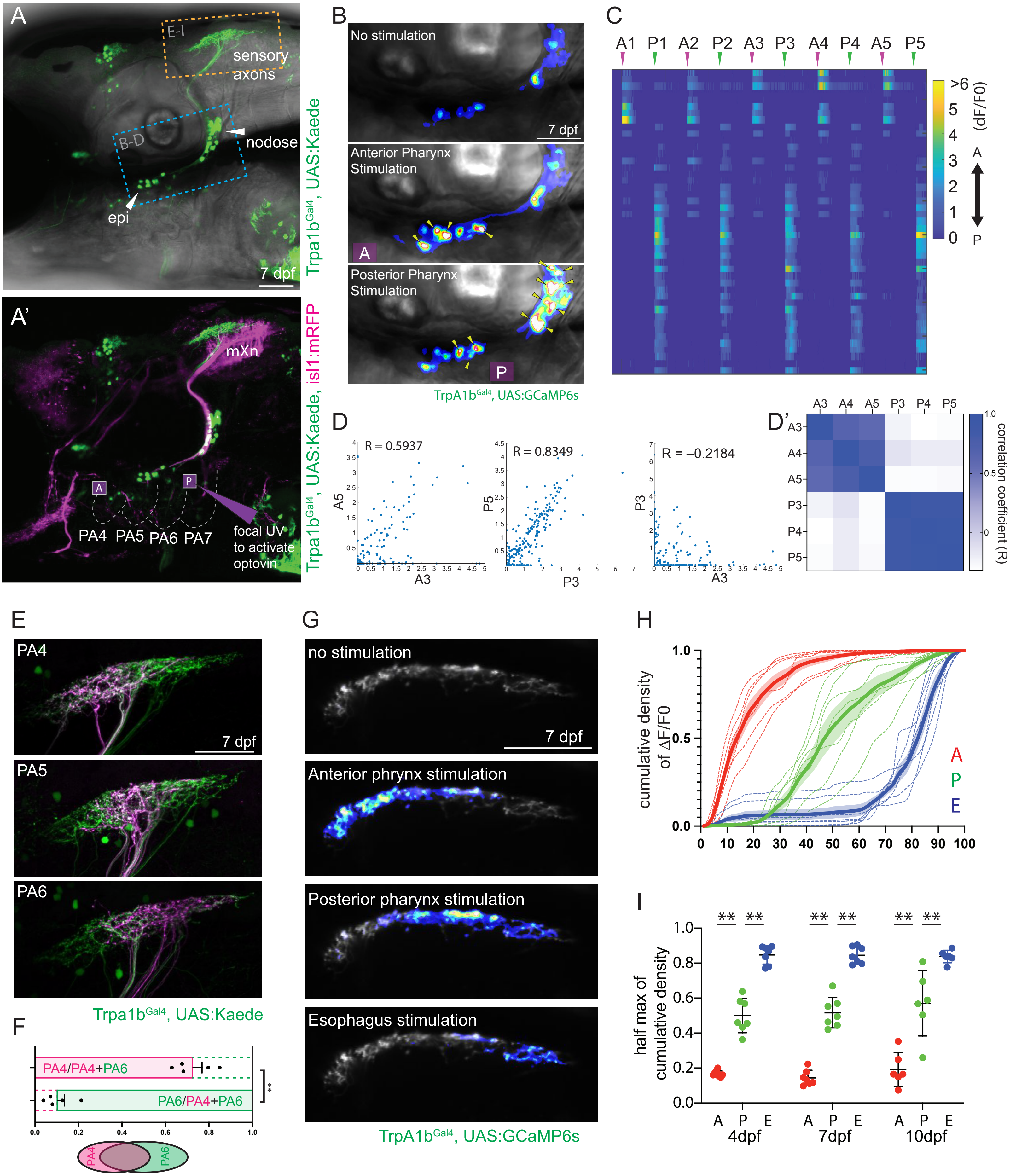

A
E-I
sensory axons
nodose
B-D
epi
7 dpf
Trpa1b$^{Gal4}$, UAS:Kaede
A'
mXn
PA4 PA5 PA6 PA7
focal UV to activate optovin
Trpa1b$^{Gal4}$, UAS:Kaede, isl1:mRFP
B
No stimulation
Anterior Pharynx Stimulation
Posterior Pharynx Stimulation
TrpA1b$^{Gal4}$, UAS:GCaMP6s
C
A1 P1 A2 P2 A3 P3 A4 P4 A5 P5
>6 (dF/F0)
D
R = 0.5937
R = 0.8349
R = –0.2184
A3
P3
A5
P5
D'
correlation coefficient (R)
E
PA4
PA5
PA6
Trpa1b$^{Gal4}$, UAS:Kaede
F
PA4/PA4+PA6
PA6/PA4+PA6
G
no stimulation
Anterior phrynx stimulation
Posterior pharynx stimulation
Esophagus stimulation
TrpA1b$^{Gal4}$, UAS:GCaMP6s
H
cumulative density of ΔF/F0
I
half max of cumulative density
4dpf
7dpf
10dpf

#### Supplemental Information

##### Figure S1 (for Figure 1). The dendrites of pharynx-innervating motor neurons are intermingled.

**(A)** Schematic of approach: Sequential photoconversion for the two pharyngeal branches, PA4 and PA6, in 4 dpf larvae expressing Tg(*isl1:Kaede*). Imaging is done 1 hour after each photoconversion, so the dendritic territory measured after the second photoconversion includes dendrites of both target groups (PA4+PA6 in this example). The square indicates the area used for quantification in **B**. **(B)** quantification of dendritic overlap between vagus target groups. The top bar shows the ratio of the PA4 dendritic area to the PA4+PA6 dendritic area, determined by sequential PA4, then PA6 (PA4>PA6) photoconversion. The second bar is the converse ratio, determined by sequential PA6>PA4 photoconversion. If there were no overlap between PA4 and PA6 dendritic territories, PA6/PA4+PA6 would be predicted to be equal to 1-PA4/PA4+PA6. In fact, PA6/PA4+PA6 (avg.=0.9, n=4) > 1-PA4/PA4+PA6 (avg 0.35, n=4), leading us to reject the null hypothesis that there is no dendritic overlap. In contrast, sequential photoconversion demonstrates no overlap between the dendritic fields of PA6 and viscera-innervating target groups (lower two bars). The level of dendritic overlap indicated from this analysis is schematized below the graph. **(C-F)** The overlay of the image after the first photoconversion (magenta) and that after the second photoconversion (green). The overlay shows the target group photoconverted first in white (i.e., magenta + green). The order of sequential photoconversion is indicated at the left. 4 dpf larvae expressing Tg(*isl1:kaede*) were used. Statistics in B: t-test. Scale bar 50 μm.

##### Figure S2 (for Figure 2). Vagus sensory neurons that express Trpa1b respond to optovin stimulation.

**(A)** Trpa1b-expressing vagus sensory neurons (green) in 7 dpf larva. Squares indicate the areas shown in **B-D** (blue) and **E-I** (orange). epi: epibranchial ganglia. **(A')** Overlay with vagus motor neurons (magenta). Purple squares indicate the areas illuminated with UV for optovin activation. mXn: vagus motor neuron. **(B)** GCaMP signals in sensory neuron cell bodies at 7 dpf following anterior and posterior pharyngeal stimulation. The magnitude of GCaMP signals is coded in the heat-color. The purple boxes indicate the region of UV illumination to activate optovin. Arrowhead indicate neurons responding to optovin activation. **(C)** Calcium traces of larva shown in **B**. Neurons are aligned from top-to-bottom corresponding to anterior-to-posterior positioning in the periphery. Noxious stimulation to the anterior (A) and posterior (P) pharynx was each repeated five times. **(D)** Correlation coefficient between activity patterns following two stimulations as in Fig. 3E. Dots represent all neurons analyzed. n= 317 neurons in 7 animals (7 dpf). **(D')** Correlation matrix for 3 iterations, with the level of correlation (R) color coded. **(E)** Anterograde labeling by Kaede photoconversion of the central processes of Trpa1b-expressing sensory neurons innervating pharyngeal arches PA4, 5, or 6. **(F)** Quantification of overlap between central processes of PA4 and PA6-innervating sensory neurons, as in Fig. S1B. **(G)** The axonal responses of vagus sensory neurons to focal optovin stimulation are topographically organized. Calcium signals following optovin stimulations are shown in the color of heat. **(H)** The anterior-posterior organization of axonal activities is quantified as cumulative density distribution of dF/F0 (see Methods) following anterior pharynx (A), posterior pharynx (P) and esophagus (E) stimulation. The A-P extent of the axon terminals is converted into the scale of 0 (anterior most) to 100 (posterior most). n=7 from 7 dpf larvae. Dotted lines show results from individual animals, with the thick line indicating the average. **(I)** Cumulative density midpoints of dF/F0 from 4 dpf (n=7), 7 dpf (n=7), and 10 dpf (n=6) are shown. Statistics in F: t-test, and statistics in I: paired t-test. Scale bars: 50 μm.

**Movie 1. UV illumination to the pharynx induces muscle contraction in the presence of optovin**

**Movie 2. UV illumination to the pharynx does not induce muscle contraction in the absence of optovin**

**Movie 3. Oil injection to the esophagus induces esophageal peristalsis through the activation of viscera-innervating motor neurons**
